# Channel width modulates the permeability of DNA origami based nuclear pore mimics

**DOI:** 10.1101/2024.05.09.593438

**Authors:** Qingzhou Feng, Martin Saladin, Chunxiang Wu, Eason Cao, Wei Zheng, Amy Zhang, Pushpanjali Bhardwaj, Xia Li, Qi Shen, Larisa E. Kapinos, Malaiyalam Mariappan, C. Patrick Lusk, Yong Xiong, Roderick Y. H. Lim, Chenxiang Lin

## Abstract

Nucleoporins (nups) in the central channel of nuclear pore complexes (NPCs) form a selective barrier that suppresses the diffusion of most macromolecules while enabling rapid transport of nuclear transport receptors (NTRs) with bound cargos. The complex molecular interactions between nups and NTRs have been thought to underlie the gatekeeping function of the NPC. Recent studies have shown considerable variation in NPC diameter but how altering NPC diameter might impact the selective barrier properties remains unclear. Here, we build DNA nanopores with programmable diameters and nup arrangement to mimic NPCs of different diameters. We use hepatitis B virus (HBV) capsids as a model for large-size cargos. We find that Nup62 proteins form a dynamic cross-channel meshwork impermeable to HBV capsids when grafted on the interior of 60-nm wide nanopores but not in 79-nm pores, where Nup62 cluster locally. Furthermore, importin-β1 substantially changes the dynamics of Nup62 assemblies and facilitates the passage of HBV capsids through NPC mimics containing Nup62 and Nup153. Our study shows the transport channel width is critical to the permeability of nup barriers and underscores the role of NTRs in dynamically remodeling nup assemblies and mediating the nuclear entry of viruses.

---

Nuclear pore complexes (NPCs) – massive (∼50–120 MDa) protein channels spanning the nuclear membranes – control molecular transport between the nucleus and cytoplasm [1, 2]. In an NPC, hundreds of nucleoporins (nups) containing unstructured Phe-Gly (FG) domains densely decorate a central channel (also known as central transporter) as narrow as ∼40 nm. These spatially confined FG-nups generate a diffusion barrier that prevents inert molecules larger than a few nanometers (∼40 kDa) from freely crossing but allows nuclear transport receptors (NTRs) to rapidly ferry cargos up to tens of nanometers in size (several MDa). The exact physicochemical state of the FG-nups in the NPC is still under debate and has been depicted as a hydrogel [3, 4], polymer brush [5, 6], and phase-separated liquid droplet [7]. Nevertheless, the multivalent and transient nup-nup and nup-NTR interactions have long been recognized as the key contributors underlying the NPC’s function as a selective barrier [8-11]. In recent years, the structural flexibility of NPC has been increasingly appreciated via in cellulo cryo-electron tomography, demonstrating central channels with diameters ranging from ∼40–70 nm depending on the cell state [12-14]. These structural studies raise the notion that the width of a nuclear pore may modulate its barrier function.

In addition to their central role in maintaining the life cycle of healthy cells, the nucleocytoplasmic transport machinery has been exploited by viruses to deliver viral genomes to the nucleus of infected cells [15]. A common strategy viruses use is to disguise their capsids—the major structural component of the virus core—as cargos for nuclear transport. To achieve this, virus capsids can incorporate nup-binding features to directly engage with NPCs (e.g., human immunodeficiency virus type 1, or HIV-1) [16-19] and/or display sequences that recruit NTRs to mediate their nuclear import (e.g., HBV) [19]. Strikingly, nearly intact capsids of certain viruses (such as HIV-1 and HBV) can cross the NPC [20, 21]. With the widths (30–65 nm) of these virus cores approaching that of the NPC’s central channel, we consider the possibility that dilated nuclear pores are preferred or even required for the transport of such ultra-large foreign objects.

The challenges to deterministically manipulate NPC structure and function have made it difficult to establish a clearcut correlation between the central channel width and NPC permeability. Due to the structural complexity of the NPC and the functional redundancy of FG-domains, there remains no obvious way to manipulate NPC diameter genetically. Structures of NPCs derived from purified nuclear envelopes are in a constricted state [22]. By contrast, biomimetic nanopores can be built with predefined geometry and nup composition while recapitulating the NPC’s basic barrier and transport functionalities [23-27]. Promisingly, a recent study showed that the transport selectivity of FG-nup-functionalized solid-state nanopores diminishes with increasing diameter beyond 55 nm [28]. However, the permeability of the NPC mimics in this study was established using relatively small proteins (<100 kDa). Furthermore, the distribution of FG domains as a function of pore diameter was not experimentally probed. In a separate study, we showed a minimal width requirement for insertion of HIV-1 capsids deep into DNA-origami-based NPC mimics (termed NuPODs for nucleoporins organized on DNA) [29]. However, such a requirement was attributed to size compatibility (i.e., a NuPOD must be at least as wide as the capsid to permit insertion) rather than altered nup behaviors in channels of different widths. Therefore, how nanopore width regulates FG-nup morphology, dynamics, and selectivity for large (>10 nm) cargos remains an open question.

To address this question, here we build NuPODs with two defined widths to mimic the central channels of constricted and dilated NPCs. Transmission electron microscopy (TEM) and high-speed atomic force microscopy (HS-AFM) reveal drastically different FG-nup morphology and dynamics in NuPODs that differ by ∼20-nm in width. The cross-channel nup meshwork and the central-plug-like Nup62/importin-β1 complexes are unique to the narrower NuPOD; the absence of these features in the wider NuPOD suggests a less stringent barrier. Corroborating these findings, the passage of HBV capsids (∼30 nm in diameter) through the narrower Nup62-Nup153 NuPOD necessitates importin-β1, an NTR known to bind both nups [30] and HBV core [19, 31], whereas the capsids can readily cross the wider pore housing the same nups without importins. The reductionist NuPOD system thus provides compelling evidence supporting a potential regulatory role for central channel width in modulating the NPC’s permeability to large-size objects, suggesting geometrical (e.g., pore diameter) and biochemical (e.g., NTR concentration) properties codetermine the barrier-transport activities of the NPC.

## Results

### Two NuPODs mimic NPC central channels of different widths

To model variations in the NPC diameter, we designed two DNA-origami nanopores that differ chiefly in width (60 nm versus 79 nm, **Figure 1A** and **1B**). Both nanopores are 33-nm deep dimers, have an octagonal cross-section, and display single-stranded DNA extensions (handles) on the interior for attaching nups. Accounting for the lengths of DNA handles (typically 21 bp or ∼7 nm long), the nup anchoring points on the opposite faces of the channel are theoretically 46 nm and 65 nm apart in the two design variants, approximating the diameters of a constricted and dilated NPC central channel, respectively. A typical DNA nanopore carries 32 handles distributed symmetrically in the middle of the channel for anchoring Nup62 and another 32 handles near one end for Nup153, mimicking these FG-nups’ stoichiometry and respective residence in the central channel and nuclear basket of a native NPC [2, 32]. The two sets of handles have orthogonal sequences, and their length, copy number, and location can be easily redesigned. We name NuPODs based on width and nup composition, for example, “60-nm Nup62 NuPOD” (designed nup copy number=32 unless specified). We further designed frustum-like DNA baskets (**Figure 1C** and **1D**) that can heterodimerize with one another and dock onto one end of the nanopores via shape complementary interfaces (**Figure 1E, 1F** and **Supplementary Figure 1**). Besides providing a visual marker for the “nucleoplasm-facing” end of the NuPODs where Nup153 is attached, the narrow opening of the basket precludes the entry of objects exceeding 30 nm in diameter, paving the way for a TEM-based HBV penetration assay.

**Figure 1:**
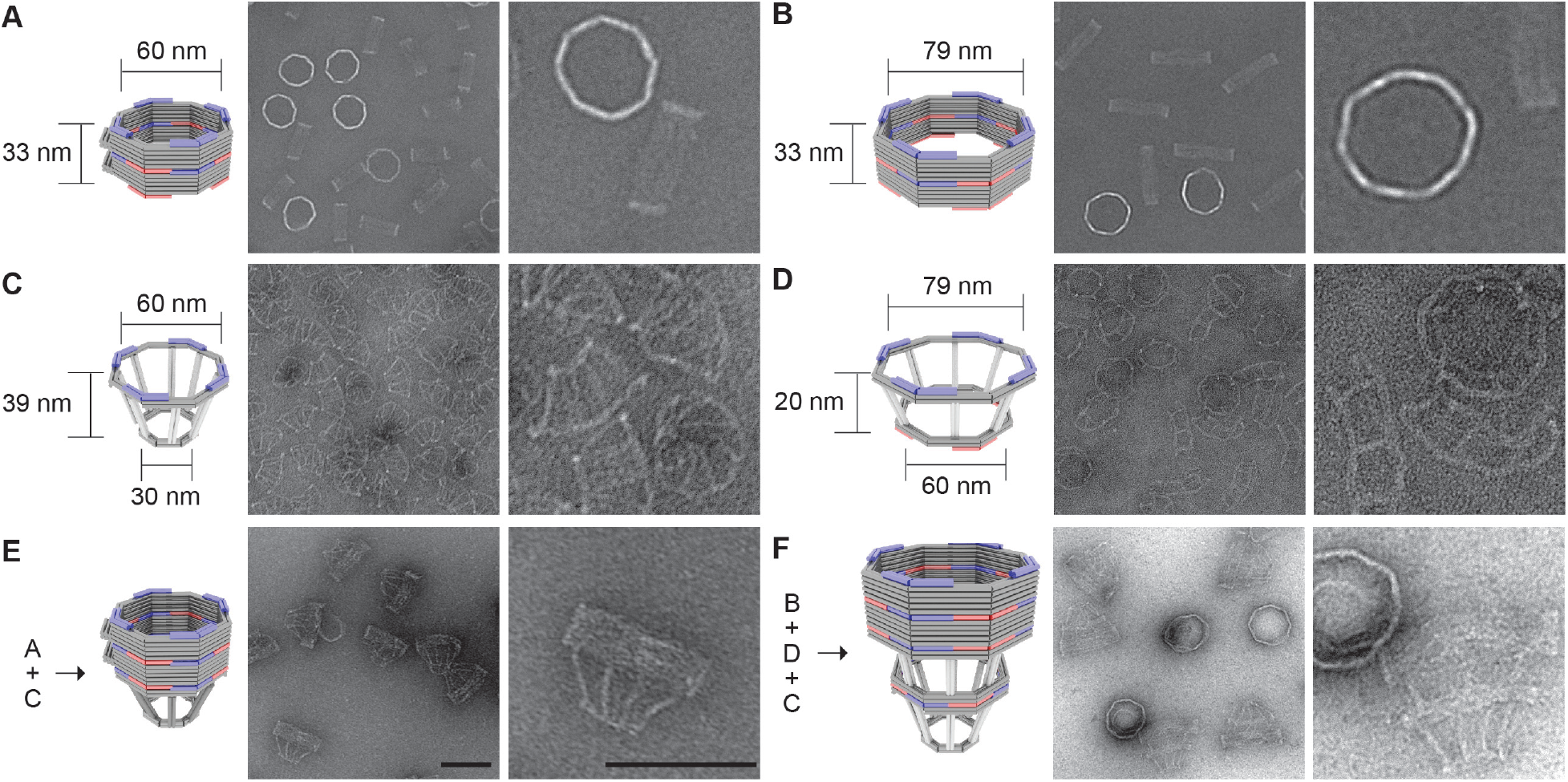
DNA-origami nanostructures used to build NuPODs. **(A)** A 60-nm wide channel built by homodimerizing DNA nanopores with shape-complementary docking interfaces (blue and red). **(B)** A 79-nm wide channel built by homodimerizing DNA nanopores with shape-complementary interfaces. **(C)** A small DNA basket with a docking interface (blue) compatible with the 60-nm channel. **(D)** A large DNA basket with top docking interface (blue) compatible with the 79-nm channel and bottom interface (red) compatible with the small basket. **(E)** A 60-nm channel capped by the small basket on one end. **(F)** A 79-nm channel capped by the two baskets on one end. For each panel, a cartoon model (DNA double helices are represented by rods) is shown next to representative TEM images. Scale bars: 100 nm.

We adapted our established workflow for assembling NuPODs [29]. All DNA-origami components (monomeric nanopores and baskets) folded with good yield and expected geometry, as confirmed by agarose gel electrophoresis and negative-stain TEM (**Figure 1C, 1D** and **Supplementary Figure 2**). Oligomerizing selected monomeric structures yielded open-ended (**Figure 1A, 1B**) or capped nanopores (**Figure 1E, 1F** and **Supplementary Figure 2**). The purified nanopores display inward facing handles that guide the subsequent nup attachment. We recombinantly expressed human Nup62^FL^ (full length, aa 1– 522) and Nup153^CTD^ (C-terminal domain, aa 896–1485) as maltose binding protein (MBP, for solubility and stability) and SNAP-tag (for DNA conjugation) fusion proteins (**Supplementary Figure 3**). Both nups conjugated efficiently with O^6^-benzylguanine-labeled DNA oligonucleotides complementary to the handles (termed anti-handles, **Figure 2A** and **Supplementary Figure 3**). Adding the purified nup-anti-handle conjugates to DNA nanopores bearing cognate handles generated NuPODs with designated channel widths and nup configurations. As the expected result of nup attachment, all NuPODs migrated slower than the corresponding empty nanopores during electrophoresis in agarose gels (**Supplementary Figure 3**). Using rate-zone ultracentrifugation to remove unbound nups and western blots to determine the amount of nanopore-attached nups, we estimated an average of ∼31 copies of Nup62 in a NuPOD, closely matching the designed number (32 copies) of Nup62 anchor points (**Supplementary Figure 4**).

**Figure 2:**
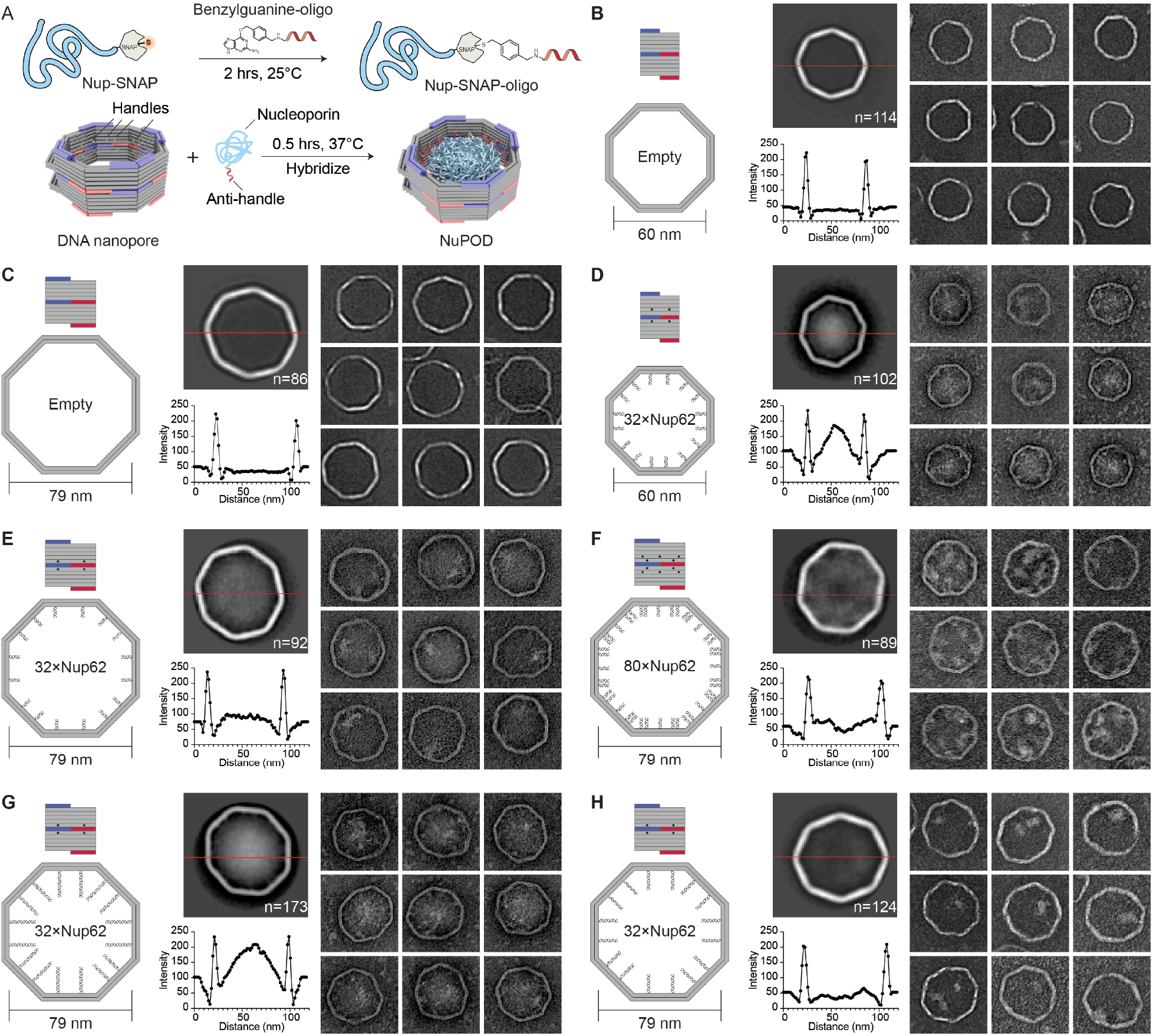
Morphology of Nup62 inside DNA-origami nanopores of different widths. **(A)** Schematic diagrams showing the process of attaching nups inside DNA nanopores to form NuPODs. **(B)** Empty 60-nm DNA channel. **(C)** Empty 79-nm DNA channel. **(D)** 60-nm NuPOD with 32 copies of Nup62 grafted on 21-nt handles. **(E)** 79-nm NuPOD with 32 copies of nup62 grafted on 21-nt handles. **(F)** 79-nm NuPOD with 80 copies of Nup62 grafted on 21-nt handles. **(G)** 79-nm NuPOD with 32 copies of Nup62 grafted on 51-nt handles. **(H)** 79-nm NuPOD with 32 copies of Nup62 grafted on 38-nt handles. For **(B)**– **(H)**, left: schematics showing an interior face (top, nup-grafting handle positions denoted by black dots) and the top view (bottom, handle/anti-handle pairs shown as double helices) of a DNA channel; middle: class average negative-stain TEM image (top) and intensity profile across the center of the DNA channel (red line); right: representative TEM images of the DNA channel. All images are 120×120 nm^2^.

### NuPOD width modulates FG-nup morphology

To probe the collective morphology of nanopore-confined FG-nups, we imaged NuPODs housing Nup62 under negative-stain TEM (**Figure 2** and **Supplementary Figure 5**). Nup62 is one of the most abundant FG-nups in the central channel of the human NPC where it plays an essential role in nuclear transport. Both 60-nm and 79-nm NuPODs were assembled without baskets to allow unobstructed view of the nups. Empty nanopores showed minimal signal in their lumens (**Figure 2B** and **2C**). Anchoring up to 32 copies of Nup62 in the 60-nm nanopore gave rise to strong protein signals (amorphous light-colored bodies against dark background) near the center of the nanopore (**Figure 2D**), suggesting the Nup62 bridged across the channel to occlude the nanopore. Such a cross-channel meshwork is missing from the 79-nm Nup62 NuPOD, where 1–2 off-center clusters formed along the channel rim (**Figure 2E**). To test whether the different morphology is due to lower nup density, which could change the conformations of intrinsically disordered domain of Nup62 from brush-like to mushroom-like, or because the polypeptide chains of Nup62 are not long enough to reach across the wider channel, we performed two sets of control experiments. We first built a 79-nm NuPOD with a maximum of 80 Nup62 proteins with higher grafting density (1/41 nm^-2^) than the 60-nm 32×Nup62 NuPOD (1/62 nm^-2^). The elevated nup density only increased the number of clusters near the rim (3–4) but did not reconstitute the pore-filling meshwork (**Figure 2F**). Notably, averaged micrographs revealed a ∼20 nm wide void in the center, suggesting very few Nup62 molecules could reach this area. Second, we used longer handles (51 bases, compared to 21 bases in typical NuPODs) to project Nup62 further inside the lumen of the 79-nm NuPODs. The additional ∼10 nm linker length appeared to help Nup62 congregate in the center (**Figure 2G**), whereas Nup62 anchored via a medium-length linker (38 base pairs) still largely formed local clusters (**Figure 2H**). The FG-domains of Nsp1 (aa 2–603), the yeast ortholog of Nup62, showed similar width-dependent morphologies in NuPODs as Nup62, albeit with a generally more diffuse appearance, perhaps because Nsp1 has weaker cohesiveness compared to Nup62 as suggested by our previous findings [29](**Supplementary Figure 6**). Therefore, we propose that the movement and interactions of nups anchored inside an NPC-like nanopore, hence their abilities to form effective pore-occluding protein bodies, are limited by nup length and the channel width as defined by the nup anchoring points.

### NuPOD width modulates the dynamics of the Nup62/importin-β1 complex

HS-AFM studies on NPCs and NPC mimics have revealed protein dynamics in the central channel on the time scale of nuclear transport events (i.e., milliseconds), which were interpreted as the hallmark of a functional barrier-transporter [33-35]. We therefore imaged NuPODs by HS-AFM to understand how pore diameter modulates nup dynamics (**Figure 3**). In keeping with our previous studies, we removed MBP tags from NuPODs by TEV cleavage to better resolve the FG-domains [25, 33, 35]. HS-AFM images and cross-sectional height profiles that were quantified from kymographs derived from rapid HS-AFM line scans (∼1.9 ms/line) showed height increases in the lumen of 60-nm Nup62 NuPODs compared with empty nanopores and further topographic elevation upon the addition of increasing concentrations (1–1000 nM) of importin-β1, confirming that the nanopore-confined Nup62 retained their native affinity to importins (**Figure 3A–3C, 3E, Supplementary Figure 7**). Remarkably, adding 100 nM importin-β1 to the 60-nm Nup62 NuPODs formed a dense yet mobile structure resembling the “central plug” in the NPC patrolling the entire nanopore [35] (**Figure 3C** and **3D, Supplementary Movie 1**). While still dynamic and capable of importin-β1 binding, Nup62 in the 79-nm NuPODs behaved very differently (**Figure 3F–3J**). The movement of Nup62/importin-β1 complexes was largely peripheral inside the 79-nm channels (**Figure 3H** and **3I, Supplementary Movie 2**), leaving a large portion of the nanopore seemingly empty most of the time. These findings are in qualitative agreement with our TEM studies and reinforce the role of channel width in regulating the collective nup behaviors.

**Figure 3:**
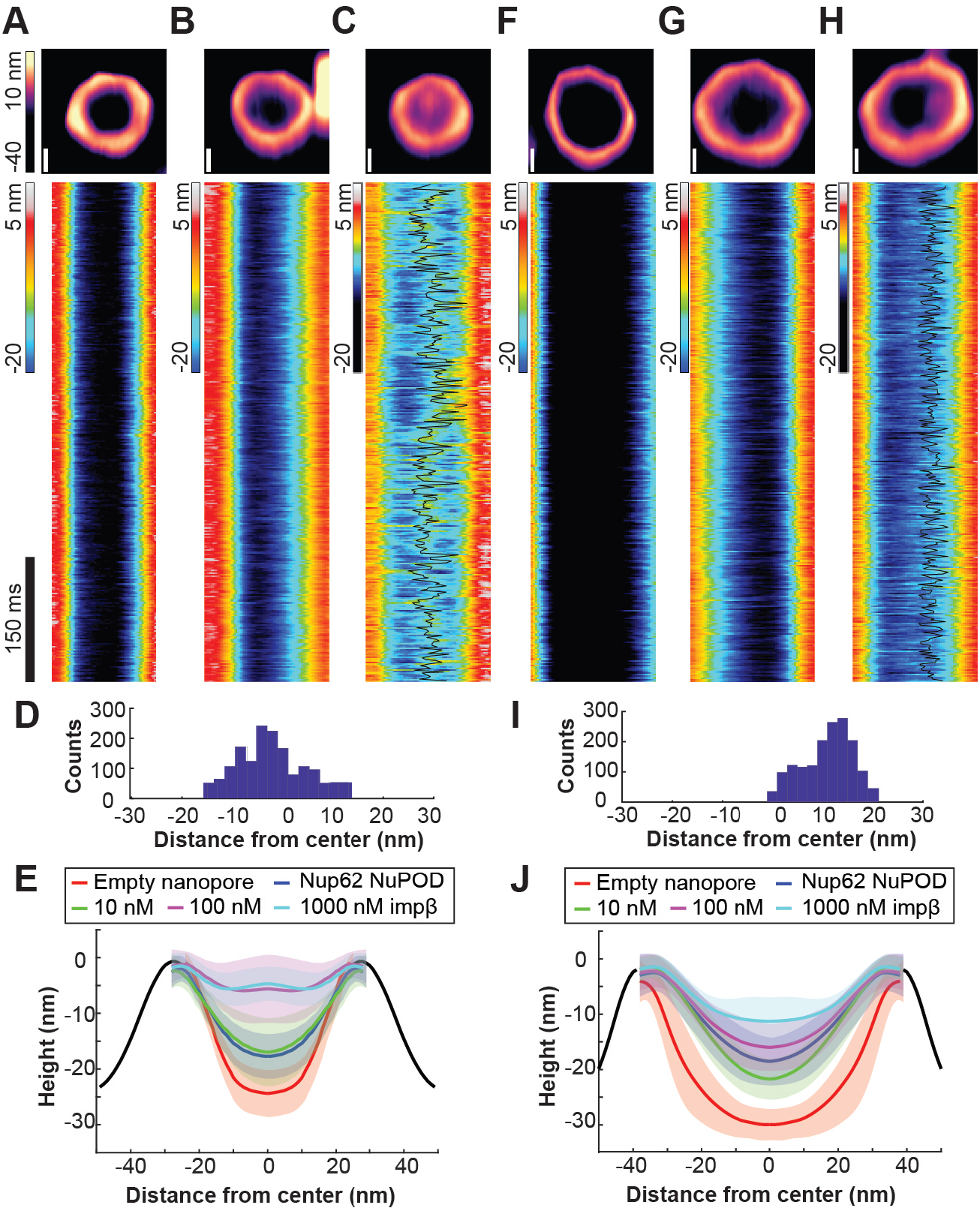
Protein dynamics inside Nup62 NuPODs of different widths. **(A)** Empty 60-nm DNA channel. **(B)** 60-nm Nup62 NuPOD. **(C)** 60-nm Nup62 NuPOD with 100 nM importin β1 (impβ). In **(A)**–**(C)** Top row: representative AFM images, scale bars: 20 nm; middle row: representative kymographs derived from HS-AFM line scans (1.875 ms/line), with movement tracking of the protein cluster overlaid (black line) for the 100 nM impβ condition. **(D)** A histogram summarizing the positions of the protein cluster in the 60-nm Nup62 NuPOD with 100 nM impβ. **(E)** Height profiles of empty nanopores and NuPODs with different concentrations of importin β1. Solid curves and shadows represent the means and standard deviations of the line scan height profiles, respectively. Number of nanopores measured: 6 (empty pore), 10 (NuPOD), 7 (NuPOD+10 nM impβ), 8 (NuPOD+100 nM impβ), and 8 (NuPOD+1 µM impβ). **(F)**–**(J)** are the same as **(A)**–**(E)**, except for the 79-nm pores. Number of nanopores measured: 12 (empty pore), 7 (NuPOD), 6 (NuPOD+10 nM impβ), 12 (NuPOD+100 nM impβ), and 11 (NuPOD+1 µM impβ).

### NuPOD width modulates Nup62 barrier permeability against HBV capsids

The absence of a central-plug-like feature that dynamically occludes the 79-nm NuPODs suggests the FG-nup barriers in the wider NuPODs may be less stringent (i.e., leakier) than their narrower counterparts. We therefore wanted to compare the permeability of 60-nm and 79-nm wide NuPODs against HBV capsids. In cells, the binding of HBV capsids with importins and Nup153, a nucleoplasm-facing FG-nup, have been proposed to mediate the translocation of the virus core through the NPC [36, 37]. We assembled HBV capsids in vitro from HBV core protein (HBc) expressed in cultured mammalian cells, generating near-spherical protein shells devoid of viral genome with an average diameter of ∼31.0±1.7 nm (**Supplementary Figure 8**). Using an ultracentrifugation-based co-migration assay (see **Methods**), we confirmed that Nup153^CTD^, but not Nup62^FL^, strongly binds HBV capsids (**Supplementary Figure 9**). Consistent with these results, HBV capsids entered basket-bearing Nup153 NuPODs efficiently, with ∼90% NuPODs occupied by least one capsid (**Figure 4A, 4B** and **4E**), while the Nup62 NuPODs showed negligible capsid binding (**Supplementary Figure 8**). Moreover, HBV capsids were largely excluded from 60-nm Nup62-Nup153 NuPODs (∼3% occupancy), suggesting that the Nup62 layer effectively blocked the Nup153-capsid interaction (**Figure 4C**). This is in stark contrast to the 79-nm Nup62-Nup153 NuPOD, whose capsid binding (∼84% occupancy) was on par with the Nup153 only NuPODs (**Figure 4D** and **4E**). From the side views of these HBV occupied NuPODs, we found a distribution of capsid penetration depths centering around the expected location of the Nup153 anchors (**Supplementary Figure 10**), reflecting Nup153-HBV binding and the extended conformation of Nup153. Because capsids must first encounter with Nup62 to engage Nup153 in the dual-nup NuPODs (the Nup153 end is capped), the increased capsid binding indicates higher permeability of the Nup62 barrier, supporting our hypothesis that wider channels lead to leakier FG-nup barriers that permit the passage of otherwise-inadmissible large objects.

**Figure 4:**
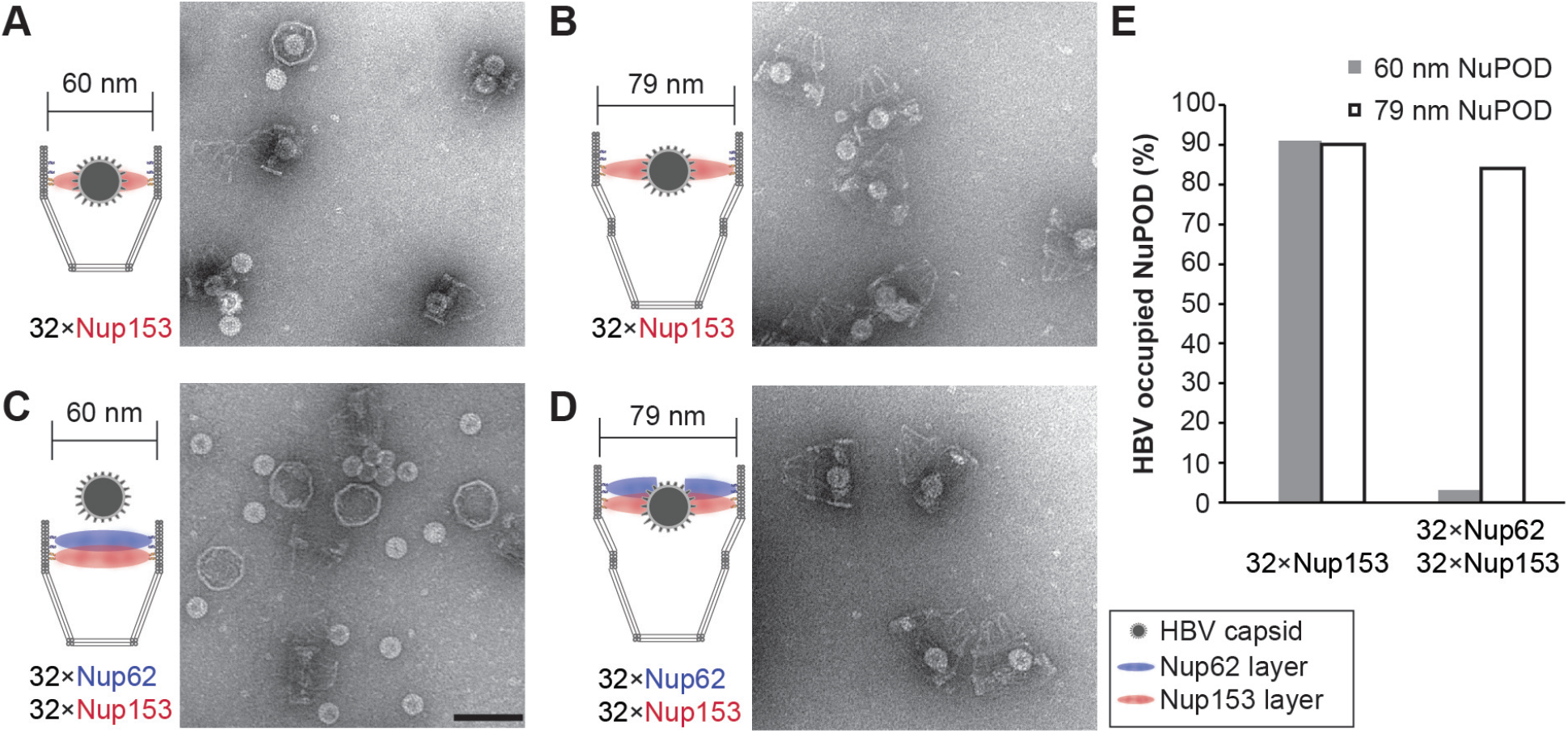
HBV capsid interactions with NuPODs of different widths. **(A)** HBV capsids mixed with capped 60-nm Nup153 NuPODs. **(B)** HBV capsids mixed with capped 79-nm Nup153 NuPODs. **(C)** HBV capsids mixed with capped 60-nm Nup62-Nup153 NuPODs. **(D)** HBV capsids mixed with capped 79-nm Nup62-Nup153 NuPODs. For **(A)**–**(D)**, schematic diagrams of the binding experiments are shown next to representative TEM images. Scale bar: 100 nm. **(E)** Percentages of NuPODs occupied by HBV capsids in experiments **(A)**–**(D)**. NuPODs counted in each experiment are (from left to right): 168, 247, 279 and 215.

### Importin-β1 mediates deep penetration of HBV capsids into NuPODs

In native NPCs, NTRs are an integral part of the selective barrier and are responsible for the transport of cellular and foreign cargos. Having established that 60-nm Nup62 NuPODs interact with importin-β1 to form central-plug-like structures, we asked whether such morphology translates to NPC-like selectivity. We continued to use the HBV capsid as a model cargo. Although nuclear import of mature HBV cores requires both importin α and importin β [38], empty HBV capsids bind directly to importin β [31], thereby simplifying our setup (**Supplementary Figure 9**). Indeed, importin-β1 substantially increased the binding between 60-nm Nup62 NuPODs and HBV capsids (**Figure 5A** and **5D, left**). Interestingly, preincubating Nup153 NuPODs with 100 nM importin-β1 resulted in a partial loss of the NuPOD’s affinity to capsids (**Figure 5B** and **5D, middle**). This is unsurprising, considering that both importin-β1 and HBV capsids compete for the Nup153 FG domain binding [39]. In the presence of importin β1, about two-thirds of 60-nm Nup62-Nup153 NuPODs were occupied by capsids, a nearly 20-fold increase compared to the importin-free condition (**Figure 5C** and **5D, right**). There appeared to be three populations of capsids in these dual-nup NuPODs, with the first two groups around the Nup62 and Nup153 layer, respectively, and the third group near the bottom of the DNA basket (**Supplementary Figure 10**). Because importin β is known to collapse Nup153 to its anchoring point [40], we interpret the deep penetration events as the result of capsids being released from Nup153, consistent with the reduced HBV binding of Nup153 NuPODs caused by importin β. In contrast to the capsids assembled from the wild type HBc, similarly sized capsids formed by HBc with a truncated C-terminus (**Supplementary Figure 8**), which bind to Nup153 but not importins (**Supplementary Figure 9**) [39], largely failed to enter the 60-nm Nup62-Nup153 NuPODs with or without importin β (**Supplementary Figure 11**). Thus, HBV capsids penetrate the Nup62 layer in the 60-nm NuPODs in an importin-dependent manner, demonstrating that these NuPODs not only form passive diffusion barriers, but also support selective, NTR-mediated transport.

**Figure 5:**
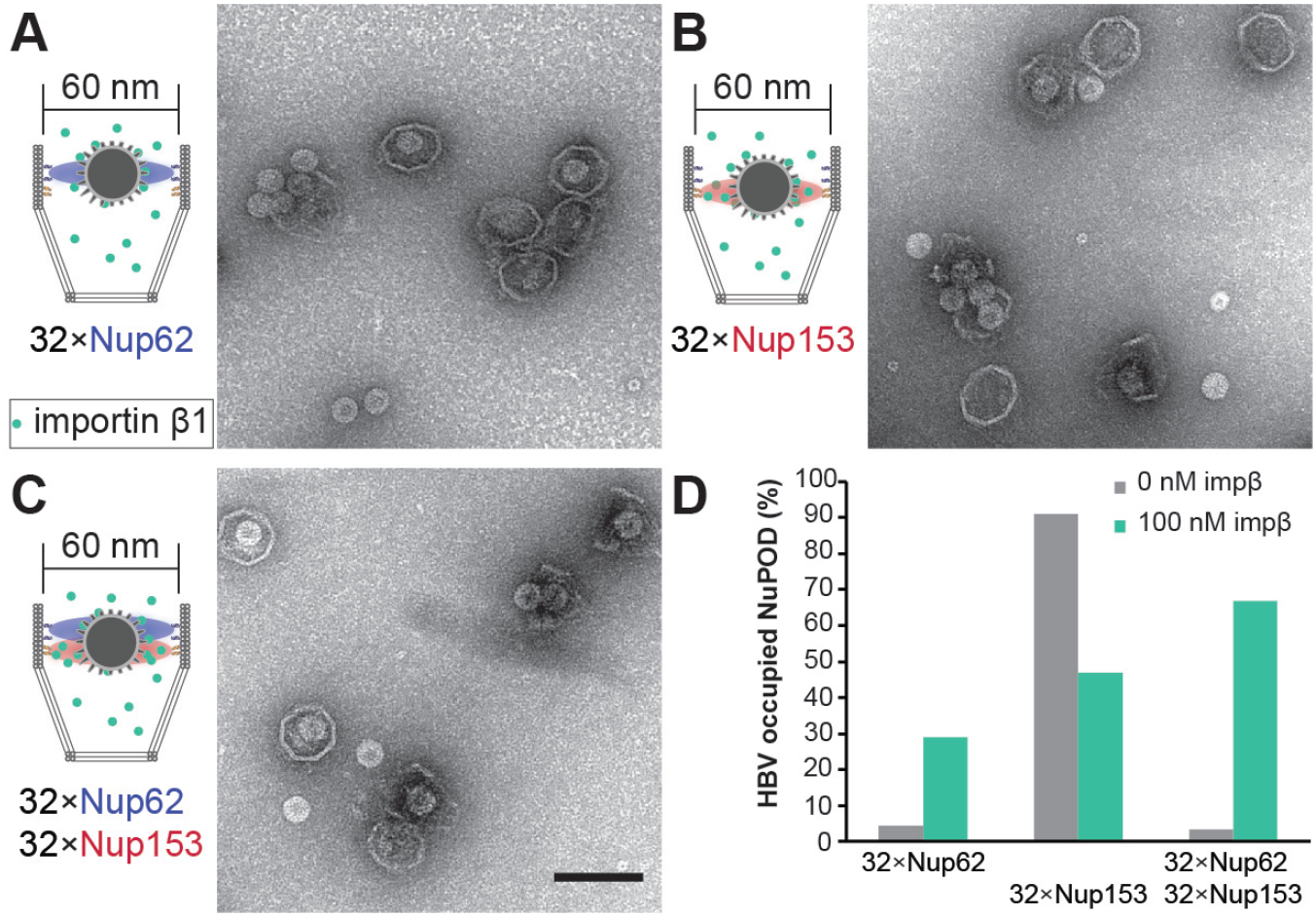
Importin mediated HBV capsid interaction with NuPODs. **(A)** HBV capsids mixed with capped 60-nm Nup62 NuPODs in the presence of 100 nM importin β1 (impβ). **(B)** HBV capsids mixed with capped 60-nm Nup153 NuPODs in the presence of 100 nM impβ. **(C)** HBV capsids mixed with capped 60-nm Nup62-Nup153 NuPODs in the presence of 100 nM impβ. For **(A)**–**(C)**, schematic diagrams of the binding experiments are shown next to representative TEM images. Scale bar: 100 nm. **(D)** Percentages of NuPODs occupied by HBV capsids with (green) or without (grey) 100 nM impβ. NuPODs counted in each experiment (from left to right): 288, 206, 168, 380, 279 and 203. Occupancies of importin-free Nup153 and Nup62-Nup153 NuPODs (**Figure 4E**) are shown here for comparison.

## Discussion

Regulation of nucleocytoplasmic transport has been attributed to biochemical determinants, such as the charge and hydrophobicity of FG-nups [16, 41, 42], NTR-nup association [10, 11, 30], and the Ran-GTP cycle [43]. Intrigued by the heterogeneous nuclear pore diameters observed in cells [12, 14, 44] and the increased nuclear import of a transcription factor caused by stretching nuclei [45], we asked how the width of the central transport channel, a geometrical attribute of the NPC, may regulate the behavior of resident FG-nups. In this study, we showed distinct Nup62 morphologies and dynamics in DNA nanopores of different widths, which we correlated to their contrasting permeability to HBV capsids. We found increasing channel width can limit the cross-channel FG-nup interactions, hinder the formation of a dynamic central cluster in the presence of importin-β1, and leave the channel less guarded (**Figure 6**). The pore-spanning FG-nup meshwork could not be rescued by increasing the Nup62 grafting density in the wider nanopore but was restored by extending the Nup62-anchoring DNA handles, confirming that the pore diameter is a fundamental determinant of cross-channel FG-nup interactions. The width-dependent morphology is shared by Nsp1 and Nup62 despite their different tendency to self-interact, suggesting nuclear pore dilation induced barrier permeabilization may be universal, though the exact behavior of the other FG-nups and their subcomplexes warrant further study.

**Figure 6:**
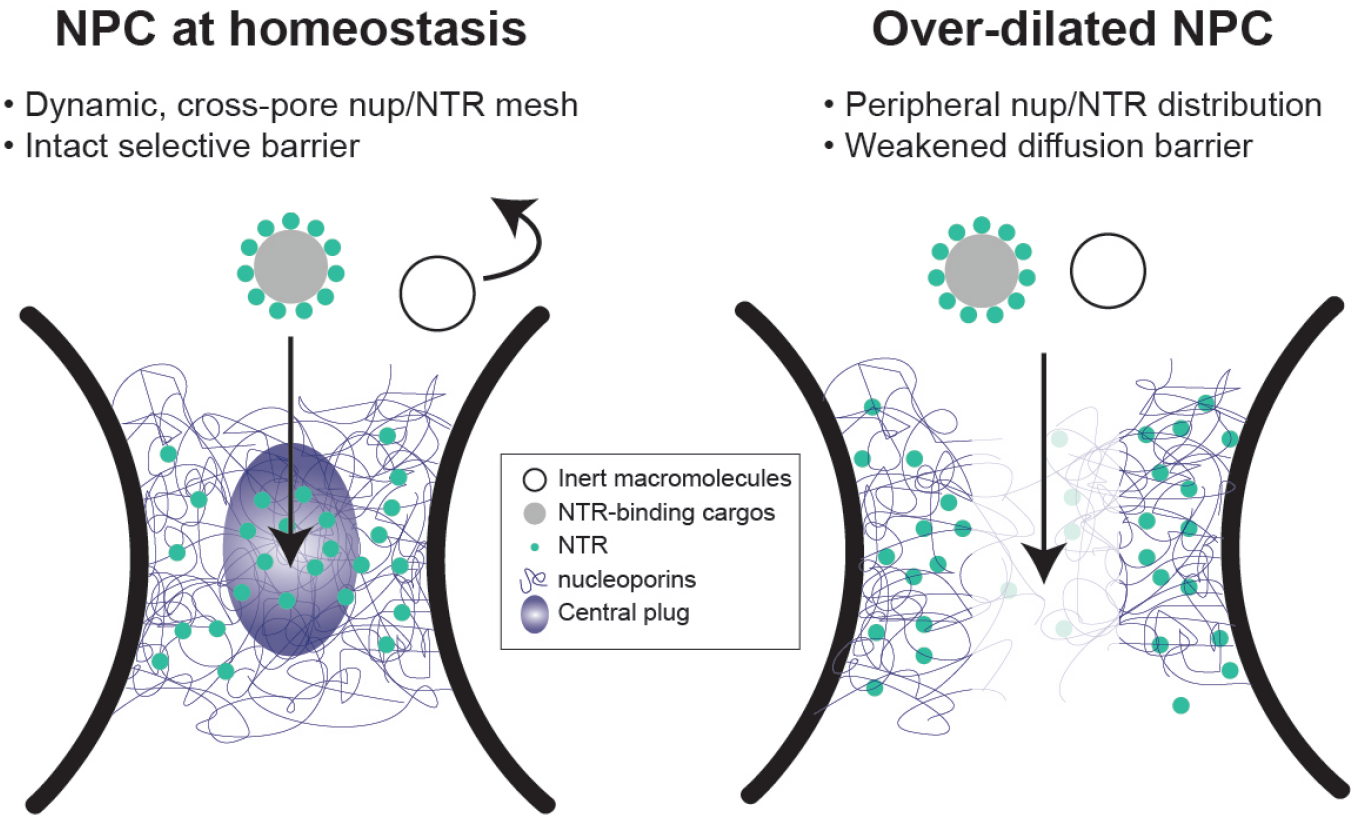
Proposed model of diameter modulated NPC selectivity. Left: Cells maintain a range of nuclear pore sizes, under which a dynamic “central plug” consisting of nucleoporins and NTRs define the selectivity of nucleocytoplasmic transport. Right: When nuclear pores dilate beyond the homeostatic range, the central channel of NPC contains less gatekeeping molecules (nups and NTRs), leading to a weaker barrier that allows non-selective transport events.

The NuPOD penetrating behaviors of HBV capsids are consistent with the previously established roles of Nup153 and importin β in mediating the HBV nuclear import [36, 37]. However, organizing FG-nups in the DNA nanopores revealed emerging properties not readily observable by canonical in vitro assays. For example, although free Nup62 showed moderate affinity toward HBV capsids, the Nup62-NuPOD almost always failed to capture capsids without the help of importin-β1. Combined with the Nup62 clustering in NuPODs, this indicates the cohesive interactions among nanopore-confined Nup62 molecules altered their binding capabilities with virus capsids, consistent with their HIV-binding behaviors in our prior work [29]. HBV capsids, which require importin-β1 to breach the Nup62 barrier in the 60-nm nanopore, went through the Nup62-decorated 79-nm nanopores independently of importin, suggesting nuclear pore dilation, in addition to NTRs, may aid the nuclear import of the virus and possibly other large objects (**Figure 6**).

Even though our current penetration assay only captures the equilibrium state (after 1 hour) of the capsid-NuPOD interaction, the TEM study still provides a glimpse of the dynamicity of the nuclear transport process. In the absence of importin-β1, interactions between NuPODs and HBV capsids were binary: nearly the entire NuPOD population was either occupied by or devoid of HBV capsids (**Figure 4**), reflecting the FG-nups’ barrier formation and HBV binding activities. Introducing importin-β1 gave rise to a range of NuPOD occupancy by capsids (**Figure 5**), suggesting the importin-associated cargos are translocated across the NuPODs with comparable on and off rates. These observations further support the role of NTRs in creating a fluid yet selective barrier, in which the transient nup-NTR interactions mediate the cargo transport [10, 30, 35, 46].

Powered by DNA nanotechnology [47, 48], the NuPOD platform has enabled the reconstitution of precisely engineered NPC mimics that form selective barriers against macromolecules [25, 27, 29, 49], affording the opportunity to dissect the sophisticated nuclear transport machinery in a reduced-complexity system. This work not only expands our toolbox by introducing more structural components (e.g., wider nanopore and baskets), but also demonstrates the value of NuPODs for the mechanistic study of nuclear import. The NuPODs complement other synthetic nuclear pore mimics, most notably the nup-functionalized solid state nanopores [23, 24, 28, 41] by allowing the precise placement of multiple FG-nup types and high-resolution study of the FG-nup-filled channel by TEM and AFM. In the future, coupling NuPODs to solid support nanopores [50, 51] or incorporating them into lipid bilayers [49, 52, 53] would enable the reconstitution of energy-dependent active transport and the measurement of molecular transport rates. Such setups would help determine the impact of transport channel geometry on the kinetics of passive and active transport. In cells, mechanical forces are thought to affect nucleocytoplasmic transport by inducing nuclear pore diameter changes and in turn leading to spatially reorganized nucleoporins [12, 14, 44, 54]. It would therefore be useful to build flexible, mechanosensitive DNA nanostructures that dynamically expand and contract to mimic the structural heterogeneity and alterations of the NPC. We expect these dynamic NuPODs to unravel mechanistic details that may have gone unnoticed in conventional cell-free experiments, such as the deformation/disintegration of virus capsids and the central channel dilation during the nuclear transport.

## Acknowledgements

C.P.L., Y. X. and C. L. acknowledge the support of NIH grant R01AI162260. R.Y.H.L. is supported by the Schweizerischer Nationalfonds zur Förderung der Wissenschaftlichen Forschung (Swiss National Science Foundation; grant no. 310030_201062). M.M. is supported by NIH grant R01GM117386.

## Author contributions

Q.F. and C.L. conceived and designed the project. Q.F., M.S., W.Z., E.C., A.Z., P.B., C.W., X.L., Q.S., and

L.E.K. performed experiments. Q.F. and M.S. analyzed data. Q.F., M.S., M.M., C.P.L., Y.X., R.Y.H.L., and C.L. interpreted data. Q.F. and C.L. prepared the manuscript. All authors participated in the discussions, and reviewed and approved the manuscript.

## Data availability

All the data are available within this paper and its Supplementary Information.

