## Supplemental Material for "Channel width modulates the permeability of DNA origami based nuclear pore mimics"

### Methods

#### Nup expression and DNA-conjugation

Genes encoding *H. sapiens* $Nup62^{FL}$ (aa 1–522) and $Nup153^{CTD}$ (aa 896–1475), both with MBP and SNAP-tag, were cloned into a pET-28a-derived vector (Novagen). To express the nups, BL21DE3 cells (New England Biolabs) were cultured at 37°C in shaking incubator until $OD_{600}$ reaches 0.5 to 1. IPTG (1 mM) was added to induce expression of target genes; the culture was kept on shaker at 20 °C overnight. The bacteria cells were then pelleted at 4,500 RPM using a Beckman Coulter JS-5.0 rotor for 15 min. Cell homogenizer (EmulsiFlex-C3, Avestin) was used to break the bacteria in a lysis buffer (50 mM Tris, pH 8.0, 150 mM NaCl, 0.1 mM PMSF and 1×RochecOmplete protease inhibitor cocktail). Cell Lysate was then centrifuge at 38,000 RPM using Beckman Coulter SW 45 Ti for 30 min to remove cell debris. The lysate containing Nup153 was then loaded to a 5 mL HisTrap column (Cytiva) on an ÄKTA system (Cytiva) and eluted using 0 to 400 mM imidazole gradient. The lysate containing Nup62 was loaded to 5 mL MBPTrap HP columns (Cytiva) and eluted by 10 mM maltose. The eluted proteins were loaded to a Superdex 200 pg 16/600 column (Cytiva) to further purify target proteins by size exclusion chromatography. The protein purity is confirmed by SDS-PAGE and Coomassie stain (Thermo Fisher).

To prepare O6-benzylguanine (BG) modified oligonucleotides (anti-handles), BG-GLA-NHS (New England Biolabs) was dissolved in DMSO for a final concentration of 20 mM. Dried 5'-amine labeled anti-handles (Integrated DNA Technologies) were resuspended in Milli-Q ultrapure water at the stock concentration of 2 mM. The amine-DNA (0.33 mM) and the NHS ester crosslinker (10 mM) were allowed to react in in HEPES (67 mM, pH 8.5) buffer for 30 min in room temperature (RT). The BG-DNA was purified by ethanol precipitation to remove unreacted crosslinkers and resuspended in Milli-Q ultrapure water. Purified BG-DNA (10 µM) was mixed with nups with a SNAP-tag protein (5 µM) in 50 mM Tris (pH 8.0) buffer and incubated at RT for 2 h. The reaction mixture was then loaded to a Superdex 200 pg 16/600 column (Cytiva) to purify the anti-handle conjugated nups from unreacted BG-DNA.

#### Importin β1 preparation

Importin β was expressed and purified according to established protocols [55], with the addition that the importin-β1-containing fraction was dialyzed against the TEV protease cleavage buffer: 10 mM Tris-HCl, pH 8, 100 mM NaCl, 1 mM DTT and 1 mM EDTA. After the $His_6$-tag cleavage by TEV protease (expressed in our lab) importin β1 was purified again using the Ni-NTA column (Roche). This was followed by additional purification using a Superdex 200 size exclusion column. The collected fraction was validated using a 12% SDS gel and flash-frozen at -80°C.

#### DNA origami design and preparation

DNA structures were designed using caDNAno [56]. DNA nanopores with handles extended from the 3' end of staple strands were shown in **Supplementary Figure 1**. The handle sequences for attaching Nup62 and Nup153 are 5′-CTACCATCTCTCCTAAACTCA-3′ and 5′-AAATTATCTACCACAACTCAC-3′, respectively. DNA origami 3D models were rendered using *Maya* (Autodesk). To fold a monomeric DNA-origami structure, a mixture of the M13mp18-derived bacteriophage circular ssDNA (8064 nt, 20 nM) and staple oligonucleotides (120 nM each) in 1×TE buffer (5 mM Tris and 1 mM EDTA, pH 8.0) with 14 mM $MgCl_2$ was annealed using a 36-h, 85–25°C protocol. The folding products were characterized by agarose gel electrophoresis (**Supplementary Figure 2**). The desired combinations of unpurified monomers were then mixed at equimolar ratio (10 nM each) and incubated at RT with adjusted magnesium concentrations for overnight to obtain dimers (14 mM $MgCl_2$), trimers (14 mM $MgCl_2$) or tetramers (24 mM $MgCl_2$). The oligomers (open-ended and capped nanopores) were purified by rate-zonal centrifugation purification as previously reported [57]. Briefly, to make the glycerol gradient, two layers of glycerol solutions (15% and 45%) containing 1×TE buffer, 14 mM $Mg^{2+}$ were load to a 13×51 mm centrifuge tube (Beckman Coulter). The quasi-linear glycerol gradient was built using a gradient station (BioComp Instruments). The unpurified DNA nanopores were loaded to the top of the gradient media and centrifuged at 45,000 RPM for 1.5 h followed by fraction collection. Fractions containing the correctly assembled nanopores (determined by agarose gel electrophoresis) were combined. Glycerol was subsequently removed by buffer exchange to 1×TE, 15 mM $MgCl_2$, pH 8.0 buffer using Amicon Ultra centrifugal filters with NMWL of 10 kDa (MilliporeSigma).

**Assembly of NuPODs**

The NuPODs were assembled from purified DNA nanopores and nup-DNA conjugates as previously described [29]. Anti-handle conjugated nups (50 mM Tris pH 8.0, 300 mM NaCl, 0.2 mM TCEP) were added to nanopores (1×TE, 15 mM $MgCl_2$) at 2:1 (anti-handle:handle) ratio. For example, the mixture for assembling 60-nm Nup62 NuPODs contained 5 nM 60-nm DNA nanopore carrying 32× handles and 320 nM anti-handle conjugated Nup62. The mixtures were incubated for 0.5–2 h at 37°C. The formation of NuPODs was validated by the retardation of NuPOD bands compared with the corresponding empty nanopores in SDS-agarose gels (**Supplementary Figure 3H**–**3J**). The same protocol was used for assembling NuPODs for the HS-AFM study with two exceptions: (1) $MgCl_2$ was adjusted to 20 mM, and (2) immediately before HS-AFM analyses, MBP was cleaved from NuPODs by incubating with TEV protease (1:50 ratio to the substrate) and DTT (1 mM) for 1.5 h at 30°C.

**Determining the nup copy number in NuPODs**

The copy number of Nup62 in NuPODs was determined using a protocol described in [25] with the results summarized in **Supplementary Figure 4**.

First, The Nup62 NuPODs were purified by rate-zonal centrifugation. Glycerol gradients were built in 5×41 mm centrifuge tubes by sequentially layering 45%, 40%, 35%, 30%, 25%, 20%, 15% glycerol solutions in 1×TE buffer containing 14 mM $MgCl_2$ (90 µL each). The Nup62 NuPODs were loaded to the top of the glycerol gradient and centrifuged at 45,000 RPM for 1.5 h. After centrifugation, nine consecutive 75 µL solution fractions were collected from top to bottom; the last fraction (<75 µL) contained aggregates. All 9 fractions were analyzed by western blot (see below). The first 8 fractions were loaded into separate lanes of an SDS-agarose gel alongside the empty nanopore as a control. The amount of NuPOD was estimated by comparing band intensities (stained by ethidium bromide) with the control sample (empty nanopore), the concentration of which was measured by $OD_{260}$. Fractions 6, 7 and 8, which contained NuPODs, were combined.

Next, the purified 60-nm Nup62 NuPOD and a series of Nup62 solutions with known concentrations (16.6–498 nM) were loaded into separate lanes of a 4–12% acrylamide gel (Thermo Fisher). Proteins were separated by SDS-PAGE, and then transferred to a nitrocellulose (Bio-Rad). Immunoblotting was performed using Nup62 polyclonal antibody (Bethyl Laboratories, Inc., A304-942A-T). Goat anti-Ribbit HRP-conjugated secondary antibody (Jackson Immunoreserach, Cat. #111-035-003) was used for detection. Band intensities on the western blot were measured using ImageJ, from which a calibration curve of band intensity vs Nup62 concentration was generated. The band intensity of the purified NuPOD in the western blot was compared to the calibration curve to determine the Nup62 concentration. The average Nup62 copy number was calculated as [Nup62]/[NuPOD].

**Negative-stain TEM and data analyses**

TEM specimens were prepared by depositing samples onto glow discharged formvar/carbon-coated copper grids (Electron Microscopy Sciences) followed by staining with 2% (w/v) uranyl formate solutions. After wicking away the staining solution, the airdried grids were imaged using a JEOL JEM-1400Plus microscope with a bottom-mount 4k×3k CCD camera (Advanced Microscopy Technologies). Particles (nanopores and NuPODs) were manually picked from TEM images (**Supplementary Figure 5**). Class averages were generated by *RELION* [58]. The intensity profiles were measured using *Fiji* [59].

**Sample preparation for HS-AFM measurements**

Supported lipid bilayers were prepared as previously described [25]. Dipalmitoylphosphatidylcholine (DPPC, Avanti Polar Lipids) and Didodecyldimethylammonium bromide (DDAB, Avanti Polar Lipids) lipids were dissolved in chloroform and mixed in a 3:1 molar ratio. The solvent was initially allowed to evaporate under a gentle nitrogen stream within a fume hood for 1 hour and then placed in a desiccator for at least 4 h. MilliQ water was added to the dry lipids to reach a final concentration of approximately 1 mg/mL. The Lipid suspension was then treated in an ultrasonic bath (Elmasonic p30H, Elma Schmidbauer GmbH) at ~65°C, using a pulse setting at 80 kHz for 15 min. The lipid solution was then loaded into syringes of an Avanti mini-extruder kit (Avanti Polar Lipids), maintaining a temperature of ~65°C, and passed through a Nuclepore Track-Etched Membrane (100 nm pore size) that is supported by a PE drain disc on both sides (Global Lifesciences Solutions UK Ltd, Buckinghamshire, UK). The process was repeated at least 20 times, resulting in the formation of small unilamellar vesicles (SUVs).

A 2.7 µL droplet of vesicle buffer (1 µL of 1 M $MgCl_2$, 1 µL of 1 M $CaCl_2$, and 16 µL of Milli-Q water) was deposited onto an HS-AFM sample stage, which consisted of a freshly cleaved mica sheet glued to a glass cylinder. SUVs (0.3 µL, described above) were then added to the droplet. The sample stage was then exposed to a temperature of ~65°C for 20 min inside a humid petri dish, followed by a gradual reduction to RT over 20 min. This thermal treatment induced vesicle rupture, leading to the formation of a positively charged supported lipid bilayer (SLB). Any excess vesicles in solution were removed by rinsing with water, followed by exchange with PB-Mg buffer (10 mM sodium phosphate, 40 mM $MgCl_2$, pH 7.0). This washing step was repeated 4 times before introducing 3 µL of TEV-treated NuPODs (2 nM). Free Nup62 and MBP in solution were removed by gently washing the stage with 1×TE buffer, 40 mM $MgCl_2$ and buffer exchange with PB-Mg buffer before HS-AFM imaging.

After depositing the NuPODs on the lipid bilayer, importin β1 was added to the sample in the imaging buffer at specified concentrations (10 nM, 100 nM, or 1 µM) and incubated for 30 min prior to the measurements.

**HS-AFM measurements on NuPODs**

High-speed atomic force microscopy (HS-AFM) measurements and data processing were performed as described elsewhere [35]. The acquisition of all HS-AFM data was conducted using the HS-AFM 1.0 system (RIBM, Japan). The system employed a standard scanner operating in tapping mode. Throughout the experiments, pristine QUANTUM-AC10-SuperSharp and QUANTUM-AC10-SuperSharp-enhanced cantilevers (nanotools GmbH) were utilized, featuring a tip radius of ⩽ 2 nm, a nominal spring constant of 0.1 N/m, a resonant frequency of approximately 0.5 MHz, and a quality factor of around 2 in water. The typical set point amplitude ($A_{set}$) was maintained at 80–90% of the free cantilever oscillation amplitude ($A_{free}$), set within the range of 2–3 nm [60, 61].

**HS-AFM line scanning measurements on NuPODs**

In high-speed atomic force microscopy line scanning (HS-AFM-LS), the slow-scan axis was deactivated, and solely the fast-scan axis was engaged [62]. A NuPOD was first imaged and centered within a region of interest prior to commencing line scanning. This ensures that the NuPOD's center is scanned repetitively, generating a kymograph as a function of the distance from the pore center. Line scans were captured over 80 pixels in the fast-scan axis at a rate of 1.875 ms per line.

**HS-AFM data processing**

All 2D images acquired through HS-AFM were drift corrected using an in-house Python-based software, which concurrently transformed the files into TIFF format [34, 35]. Subsequent analysis of HS-AFM images was done using ImageJ along with custom analysis routines written in Python. Various image processing tasks, such as filtering, contrast adjustment, and height/diameter measurements, were also performed using ImageJ. Initially, all HS-AFM images underwent correction for XY-tilting through the application of a flattening filter utilizing a first-order polynomial plane. Following this, the heights of the HS-AFM images were normalized in relation to the scaffold by subtracting the average scaffold height. Zoomed-in HS-AFM images were treated using a 2D Gaussian filter with a 1-pixel standard deviation.

HS-AFM-LS data was processed using custom analysis routines written in Python as follows. Full kymographs were spliced into 600 ms segments. Corrections for drift in the Z-axis and tilt in the fast-scan axis were applied, and the heights were adjusted relative to the NuPOD scaffold, provided the top surface of the scaffold was discernible within the same kymograph. Furthermore, a Fast Fourier transform filter was applied to eliminate periodic noise at frequencies of 50 Hz and 150 Hz, with root mean square (RMS) amplitudes of approximately 0.09 nm and 0.08 nm, respectively. Given their negligible size, these amplitudes had no discernible impact on the dynamic analysis of HS-AFM-LS. The height values obtained from kymographs were averaged along the time axis at corresponding radii and then overlaid over the average profile of the scaffold (**Figure 3E** and **3J**).

**Particle-tracking and positional distribution**

A custom Python routine tracked the displacement of the central plug-like feature in HS-AFM kymographs on a line-by-line basis along the fast-scan (X) axis [35]. Initially, a low-pass filter was applied to the kymographs along this axis to enhance the signal-to-noise ratio. Detection was specifically limited to the central region, excluding the NuPOD scaffold. If displacement of the CP-like cluster was detected within a line, the algorithm selected the position of the highest local maximum; otherwise, the line was skipped. Trajectories from all lines containing the cluster were then compiled into a histogram to display the distribution of the cluster relative to the center of a NuPOD (referenced in Figure 3D and 3I).

**In vitro assembly of HBV capsids of mammalian cell origin**

Expi293 cells were transfected with wt HBV core protein (HBc) cloned in the pcDNA3.4 mammalian vector (Thermo Fisher Scientific). After culturing at 37°C for 72 h, the cells were harvested. The cell pellet was resuspended in a lysis buffer (50 mM Tris-HCl pH 8.5; 150 mM NaCl; 1% NP-40; protease inhibitors) and chilled at 4°C for approximately 1 h. In cases of incomplete lysis, cells were manually disrupted using a Dounce homogenizer until complete lysis was observed under a light microscope.

The cell lysate was then centrifuged at 17,000 g for 45 min at 4°C. The supernatant was collected and loaded onto a 20–60% glycerol gradient (20 mM Tris pH 8.0, 150 mM NaCl). This gradient was centrifuged at 4°C for approximately 5 h at 153,700 g in a SW 32 Ti rotor. After spinning for 5 h, the entire gradient was fractionated into 1.5 mL aliquots, which were analyzed using 4–12% SDS-PAGE. After buffer exchange to remove glycerol, the fractions with the most abundant full-length HBc were imaged by negative-stain TEM. Based on TEM analysis, the fractions containing the most intact HBV capsids were concentrated and stored at -80°C before being used for binding and penetration assays. The absorption (220–350 nm, with 1 nm increment) spectrum of the capsids was measured using a NanoDrop spectrometer (Thermo Fisher).

**In vitro assembly of HBV capsids of bacterial origin**

Plasmids encoding HBc (wt or ΔC) were transfected to BL21(DE3) cells and expressed as described in **Nup purification and DNA conjugation**. After lysing the bacteria in a buffer (50 mM Tris, pH 8, 300 mM NaCl), bacterial debris were removed by centrifugation at 12,000 g for 1 h. HBc proteins were pelleted from the supernatant by adding 30% of $(NH_4)_2SO_4$ and centrifuging for 1 h at 12,000 g. The pellets were resuspended in 50 mM Tris-HCl (pH 8.0) before dialyzed overnight against a buffer (50 mM Tris, pH8). After centrifugation at 12,000 g for 1 h to remove aggregates, the dialyzed sample was loaded to a HiPrep™ Q FF 16/10 anion exchange chromatography column (Cytiva) and the flow though from the column, which contained HBc, was collected. The flow through was subjected to rate-zonal centrifugation at 30,000 rpm (SW 32 Ti rotor), 4°C for 3 h in a 20–60% glycerol gradient (20 mM Tris pH 8.0, 150 mM NaCl). Fractions collected from the gradient were analyzed by SDS-PAGE. Fractions containing HBc of the expected M.W. were collected and concentrated by Amicon Ultra centrifugal filters with NMWL of 100 kDa. The fractions with the most abundant HBc were imaged by negative-stain TEM to screen for intact HBV capsids. The capsids were stored at -80°C before being used for binding and penetration assays. The absorption (220–350 nm, with 1 nm increment) spectra of the capsids were measured using a NanoDrop spectrometer (Thermo Fisher).

**HBV penetration assay**

Purified HBVs (50 μg/ml HBc) and NuPODs (5 nM) were mixed to achieve approximately 1:1 capsid:NuPOD molar ratio and incubated at RT for 15 min. To test the effect of NTRs on HBV-NuPOD interactions, NuPODs were pre-incubated with 100 nM importin-β1 for 15 min before the addition of HBV capsids. The mixtures were then imaged by negative-stain TEM. Electron micrographs were analyzed by ImageJ to determine the HBV occupancy (number of HBV-bound NuPODs/total number of NuPODs) and penetration depth (defined as the distance from the NuPOD entrance to the center of the capsid). Practically, the penetration depth was calculated by subtracting the average radius of the HBV capsids (15 nm) from the distance measured from the NuPOD entrance to the bottom of the HBVs. All NuPOD-bound capsids were measured as long as the capsid center passed the entrance of the NuPOD. The penetration depth data were plotted as normalized histograms and fitted by multiple normal distributions using MATLAB (MathWorks).

**Comigration-based binding assays**

HBV wt (3 µM HBc) and HBV ΔC (3 µM HBc) were incubated separately with ~ 1 µM of Nup62, Nup153, or importin β1 at RT for 15 min. The mixtures were each loaded onto a linear glycerol gradient (15% to 45%, in 1×TE, pH 8, 15 mM $MgCl_2$) and spun for 90 min at 48,000 RPM using SW 55 Ti Rotor. The collected fractions (12–13 fractions in total, 50 µL each) were characterized by 4–12% SDS-PAGE to determine the co-migrating proteins (**Supplementary Figure 9**). HBV capsids, nups, and importin β1 were individually subjected to the same procedures as controls.

**Data analysis and statistics**

Scatter plots are presented as mean$\pm$ SD using the software Prism8.0.0 (Graph-Pad). The two-tailed T-test were used to compare the diameters of the HBV wt and HBV ΔC assuming normal distribution with equal variances.

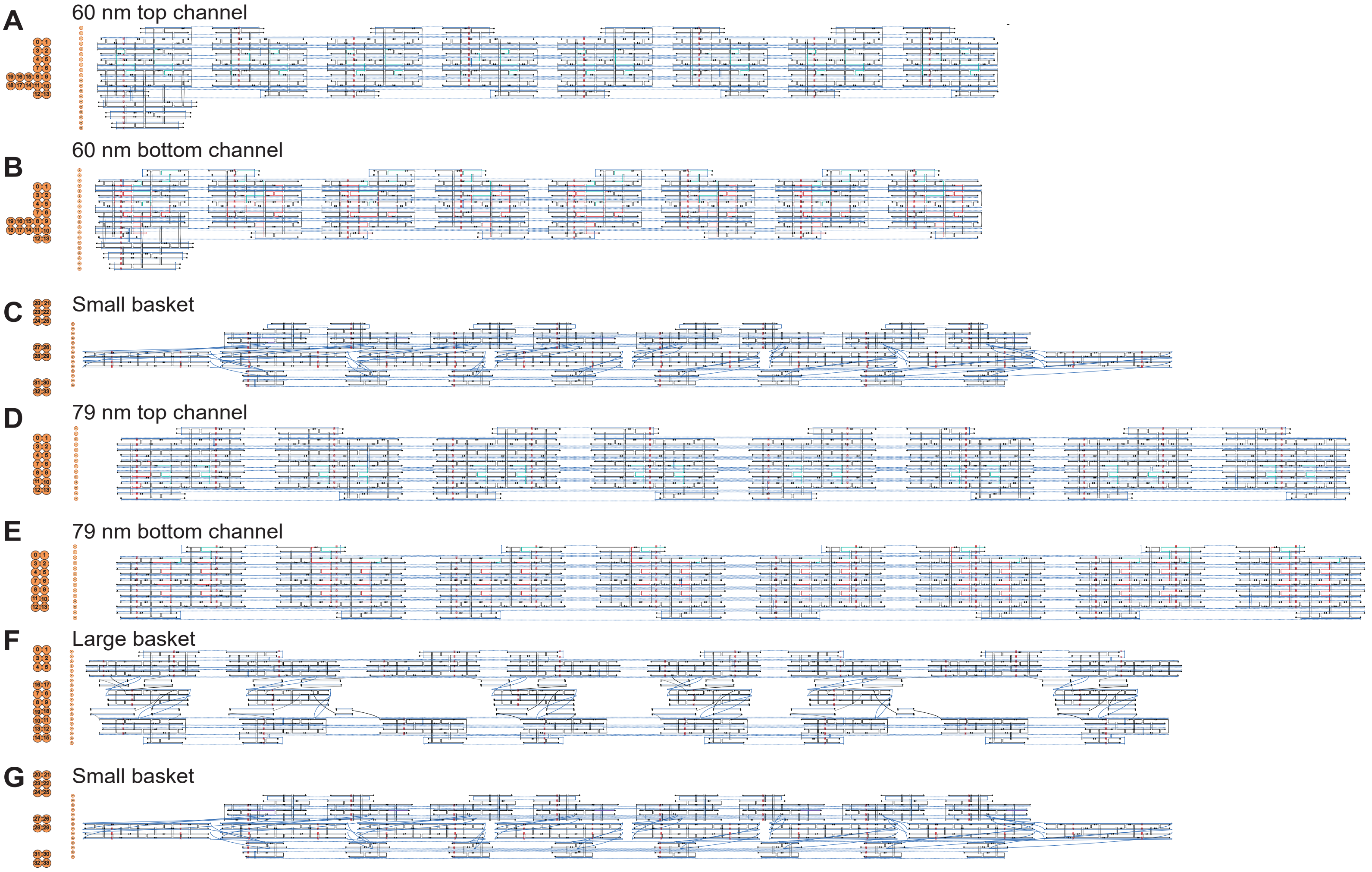


**Supplementary Figure 1:** DNA origami designs shown as caDNAno diagrams.
The 3'-ends of the green and red staple strands represent handle positions for Nup62 and Nup153, respectively.

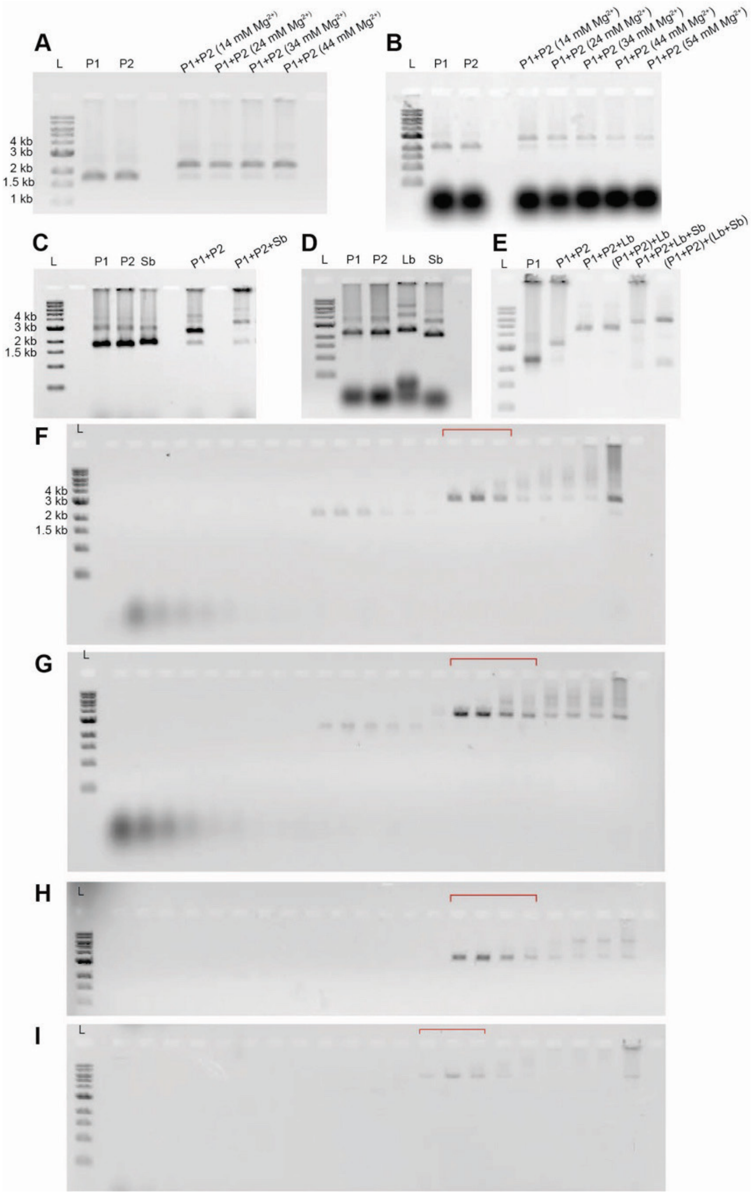


**Supplementary Figure 2**: DNA origami assembly and purification.

**(A)** Assembly of the open-ended 60-nm channel at various $Mg^{2+}$ concentrations characterized by agarose gel electrophoresis. **(B)** Assembly of the open-ended 79-nm channel at various $Mg^{2+}$ concentrations characterized by agarose gel electrophoresis. **(C)** Assembly of the capped 60-nm channel at various $Mg^{2+}$ concentrations characterized by agarose gel electrophoresis. **(D)** The 79-nm NuPOD components characterized by agarose gel electrophoresis. **(E)** Assembly of the capped 79-nm channel characterized by agarose gel electrophoresis. L: 1kb DNA ladder; P1: top channel; P2: bottom channel; Lb: large basket; Sb: small basket. Parentheses denote pre-assembled components. **(F)** Open-ended 60-nm channel purified by rate-zonal centrifugation. **(G)** Open-ended 79-nm channel purified by rate-zonal centrifugation. **(H)** Capped 60-nm channel purified by rate-zonal centrifugation. **(I)** Capped 79-nm channel purified by rate-zonal centrifugation. For **(F)–(I),** fractions 5–27 are resolved by agarose gel electrophoresis (lighter fractions on the left), and the collected fractions containing the correctly assembled structures are denoted by red brackets.

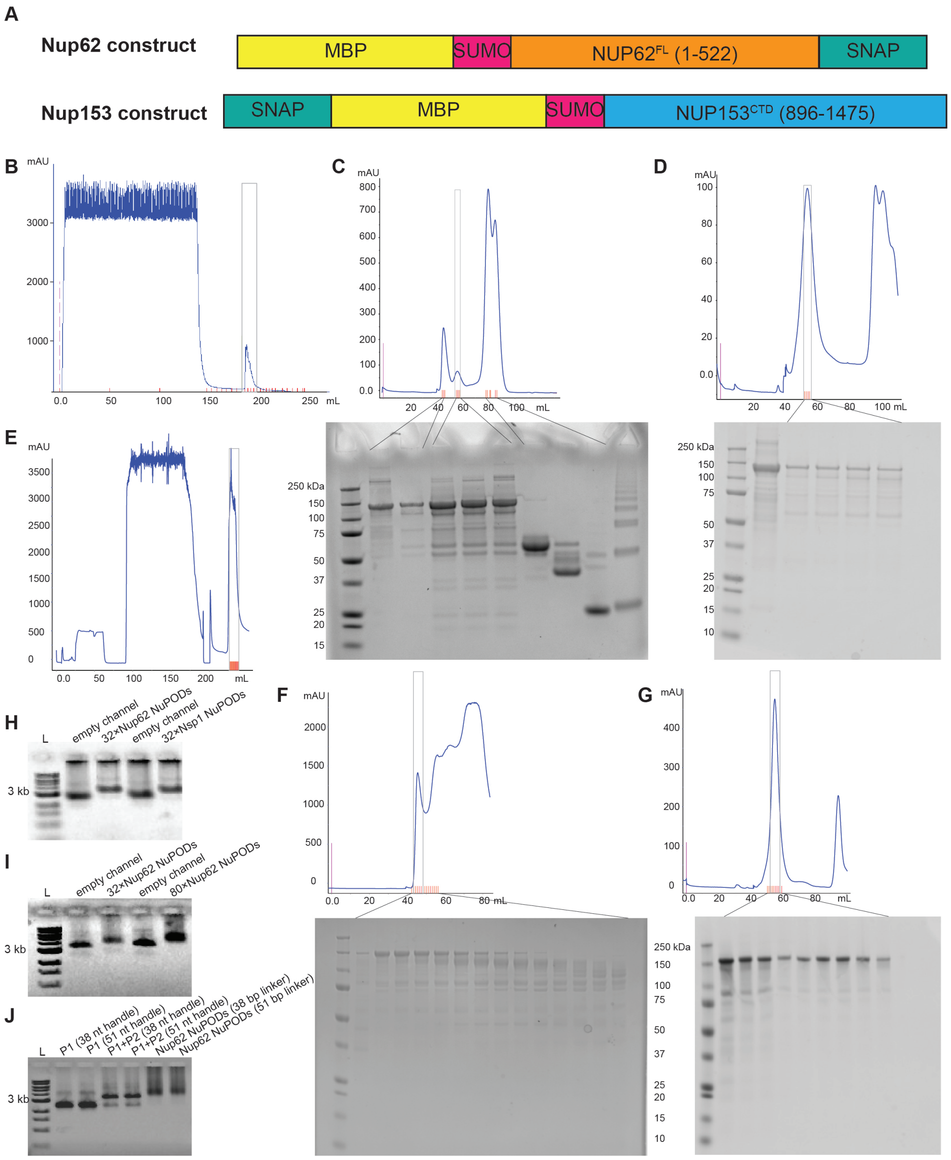


**Supplementary Figure 3**: Preparation of nups, nup-DNA conjugates, and NuPODs.

**(A)** Schematics of the Nup62 and Nup153 constructs used in this work. **(B)** Purification of Nup62 by MBP affinity chromatography. Elusions in the grey box are collected for further purification by size-exclusion chromatography (SEC). **(C)** SEC purification of Nup62. Boxed fractions are collected for DNA conjugation. **(D)** SEC purification of DNA-conjugated Nup62. Boxed fractions are collected for NuPOD assembly. **(E)** Purification of Nup153 by His affinity chromatography. Elusions in the grey box are collected for further purification by size-exclusion chromatography (SEC). **(F)** SEC purification of Nup153. Boxed fractions are collected for DNA conjugation. **(G)** SEC purification of DNA-conjugated Nup153. Boxed fractions are collected for NuPOD assembly. Selected fractions in **(C), (D), (F),** and **(G)** are characterized by SDS-PAGE. **(H)** Agarose gel (0.05% SDS) electrophoresis (SDS-AGE) showing the band shifts of open-ended 60-nm Nup62 and Nsp1 NuPODs compared with empty DNA channel dimers. **(I)** SDS-AGE showing the band shifts of open-ended 79-nm NuPODs with different Nup62 copy numbers compared with empty DNA channel dimers. **(J)** SDS-AGE gel showing the band shifts of 79-nm Nup62 NuPODs with different handle lengths compared with empty DNA channel monomers (P1) and dimers (P1+P2).

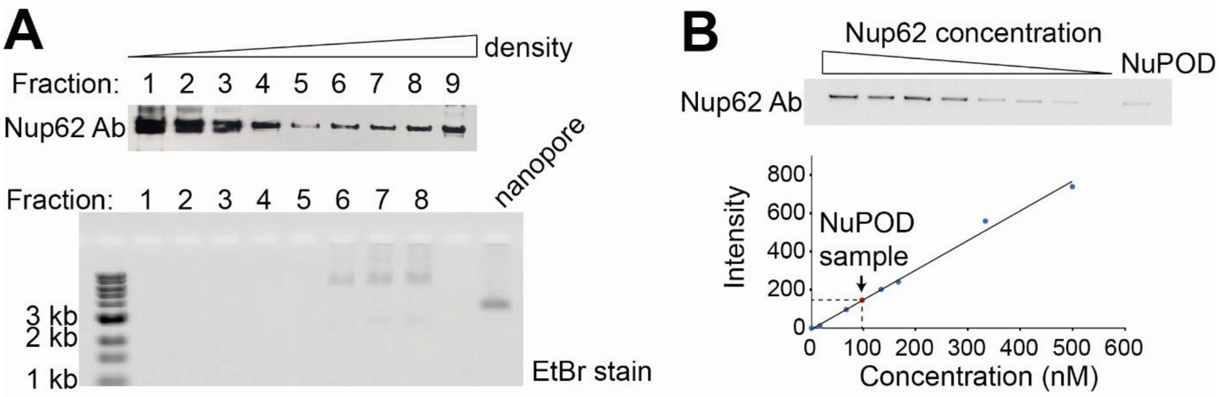


**Supplementary Figure 4**: Measuring the Nup62 copy number inside a NuPOD.

**(A)** Rate-zonal centrifugation separates free Nup62 from NuPODs. Top: Fractions recovered after rate-zonal centrifugation of unpurified Nup62 NuPODs, characterized by SDS-PAGE/western blot (detected using Nup62 antibody); Bottom: The first 8 fractions characterized by SDS-agarose gel electrophoresis (SAS-AGE) and stained by ethidium bromide. Lighter fractions are shown on the left. **(B)** Western blot for measuring Nup62 concentration in the purified NuPOD, a mixture of fractions 6–8 shown in **(A)**. Top, SDS-PAGE/western blot (detected using Nup62 antibody) of purified Nup62 (the first 8 lanes) and Nup62 NuPOD (the right most lane). Bottom: A plot showing western blot band intensity (y) against known Nup62 concentration (x), with linear regression (solid line) yielding y=1.547x-3.604 ($R^2$=0.99). The red dot represents the NuPOD band intensity and the corresponding Nup62 concentration (94.3 nM). Given the NuPOD concentration (3.05 nM, estimated by band intensity comparison with empty nanopore in SDS-AGE), we estimated ~31 copies of Nup62 in one NuPOD on average.

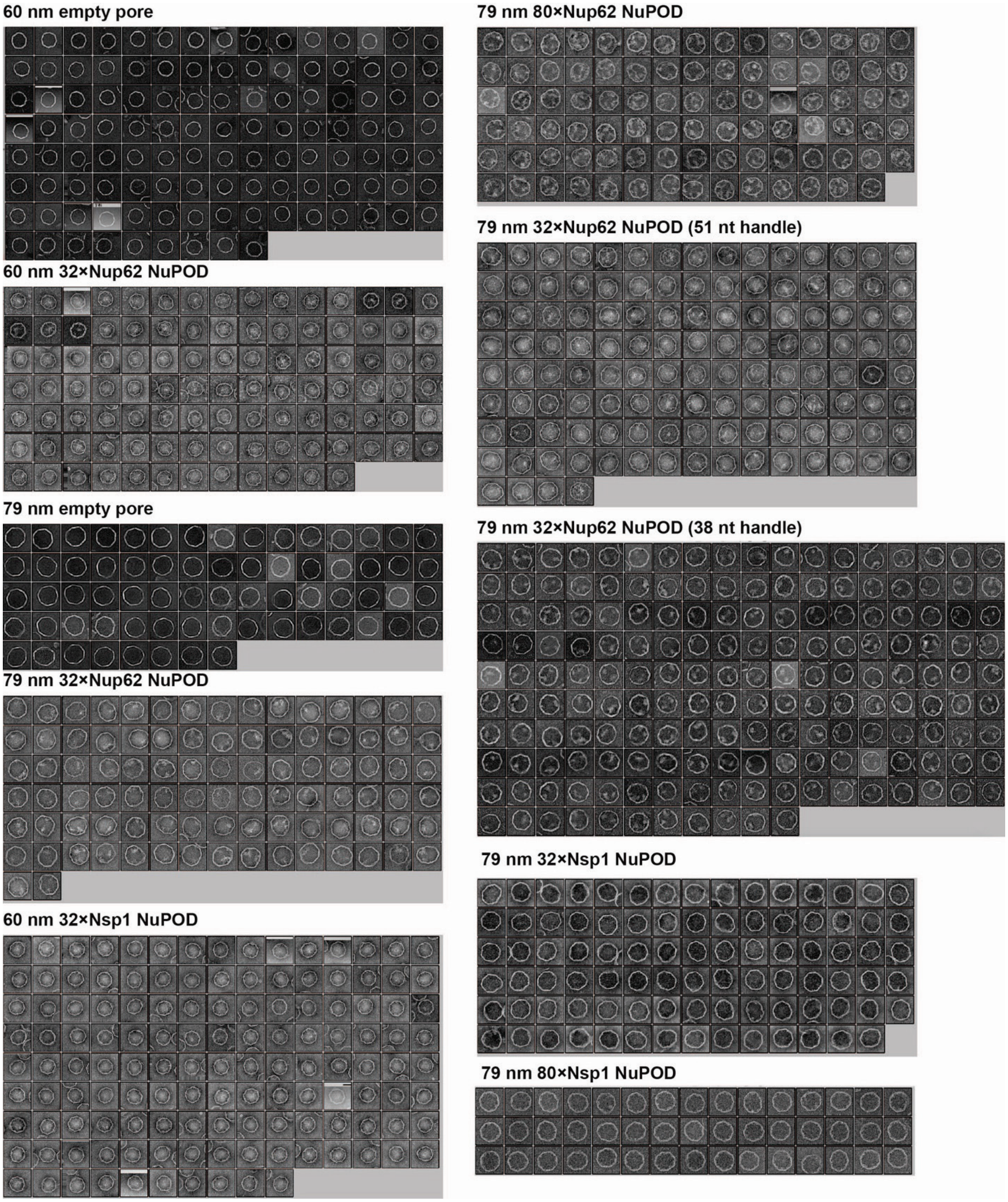


**Supplementary Figure 5**: Negative-stain TEM image galleries of empty DNA nanopores and NuPODs. The RELION 2D average and a few representative images of each sample are shown in **Figure 2** and **Supplementary Figure 6**.

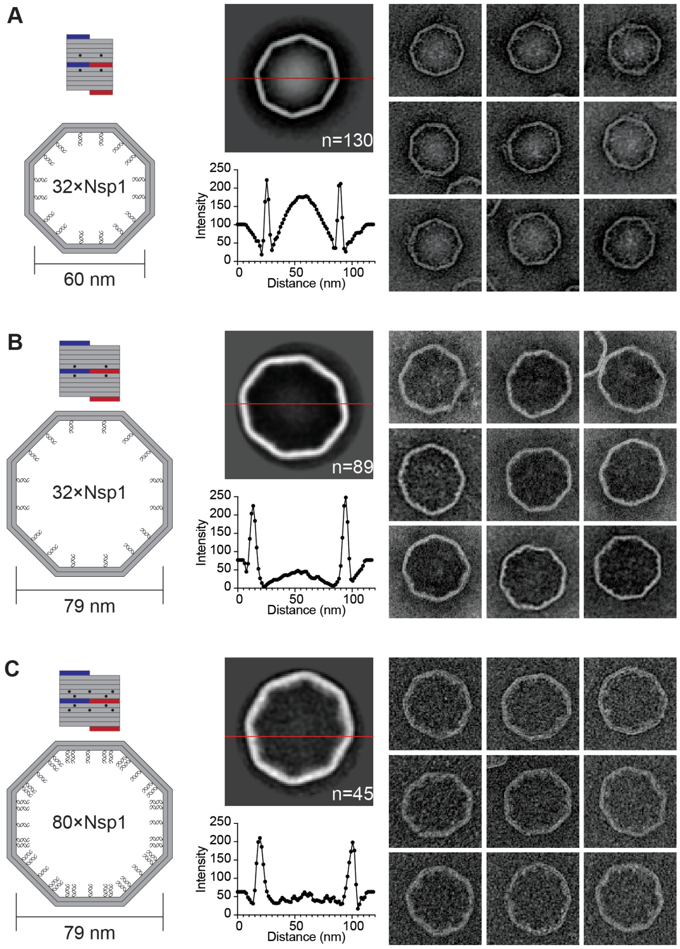


**Supplementary Figure 6**: Morphology of Nsp1 inside NuPODs of different widths.

**(A)** 60-nm NuPOD with 32 copies of Nsp1. **(B)** 79-nm NuPOD with 32 copies of Nsp1. **(C)** 79-nm NuPOD with 80 copies of Nsp1. For each panel, left: schematics showing an interior face (top, nup-grafting handle positions denoted by black dots) and the top view (bottom, handle/anti-handle pairs shown as double helices) of a DNA channel; middle: class average negative-stain TEM image (top) and intensity profile across the center of the DNA channel (red line); right: representative TEM images of the DNA channel. All images are 120×120 $nm^2$.

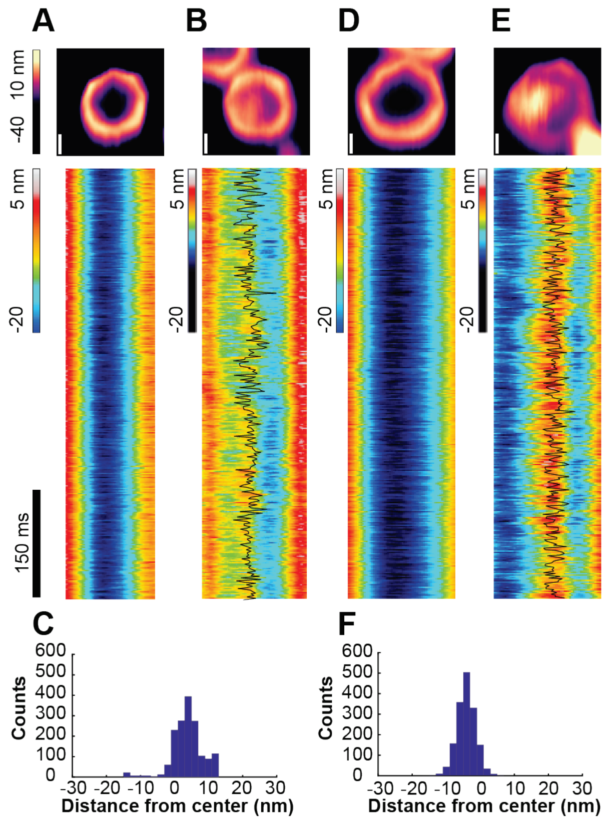


**Supplementary Figure 7**: Protein dynamics inside Nup62 NuPODs with importin β1.
**(A)** 60-nm Nup62 NuPOD with 10 nM importin β1 (impβ). **(B)** 60-nm Nup62 NuPOD with 1 μM impβ. Top row: representative AFM images, scale bars: 20 nm; middle row: representative kymographs derived from HS-AFM line scans (1.875 ms/line), with movement tracking of the protein cluster overlaid (black line) for the 1 μM impβ condition. **(C)** A histogram summarizing the positions of the protein cluster in the 60-nm Nup62 NuPOD with 1 μM impβ. **(D)**–**(F)** are the same as **(A)**–**(C)**, except for the 79-nm NuPODs.

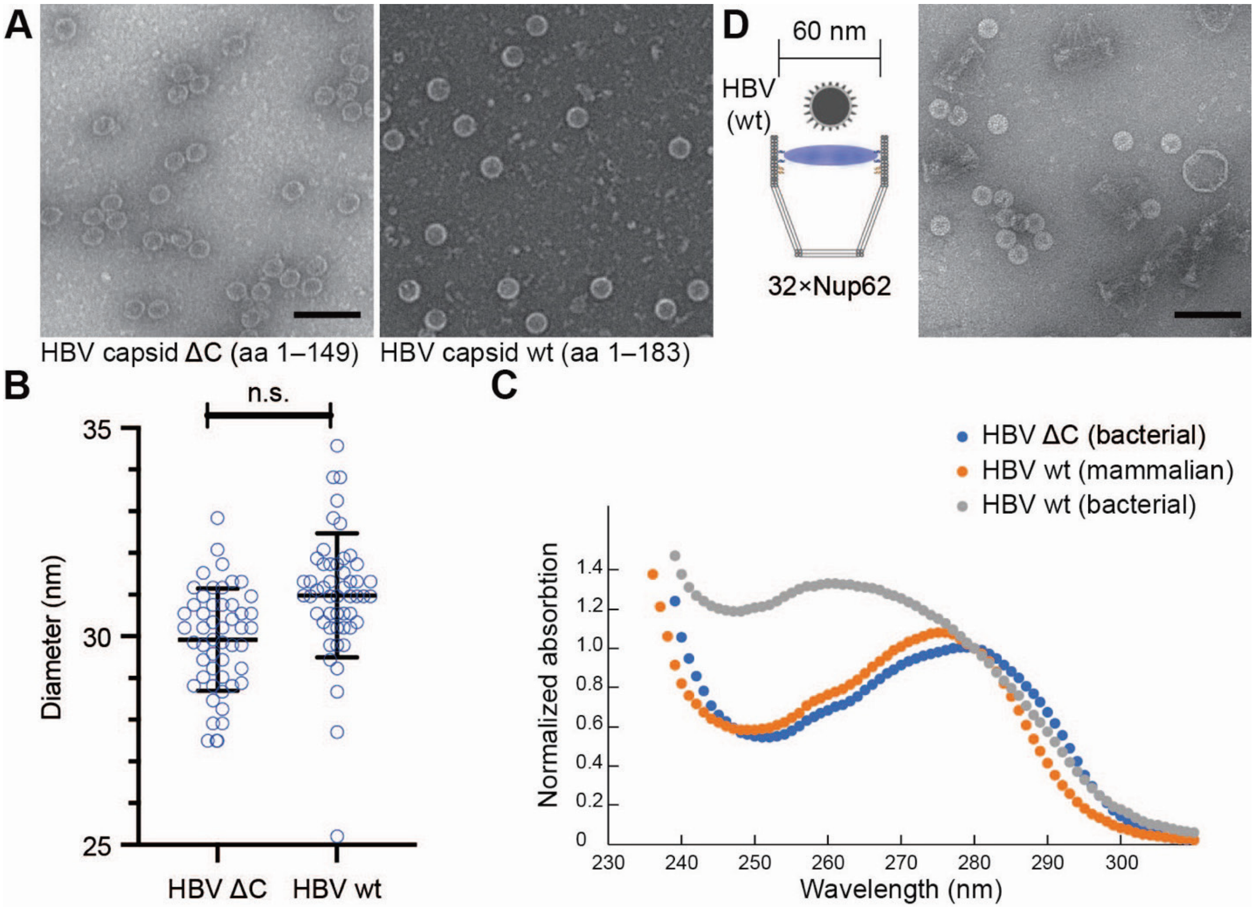


**Supplementary Figure 8**: Characterization of HBV capsids.

**(A)** TEM images of capsids assembled from wild-type (wt) and C-terminus truncated (ΔC) HBc. Scale bar: 100 nm. **(B)** Measured diameters of HBV capsid wt (30.98 $\pm$ 1.73 nm, n=50) and ΔC mutant (29.92 $\pm$ 1.11 nm, n=50). P=0.1763 based on a two-tailed t-test; n.s.: not significant. **(C)** Absorption curves of HBV capsids normalized to $OD_{280}$. Nucleic acids in capsids assembled from E. coli expressed wt HBc are indicated by large $OD_{260}$ (grey line). **(D)** HBV capsids (wt) mixed with 60-nm Nup62 NuPODs capped at one end. A schematic of the binding experiment is shown next to a representative TEM image. Scale bar: 100 nm.

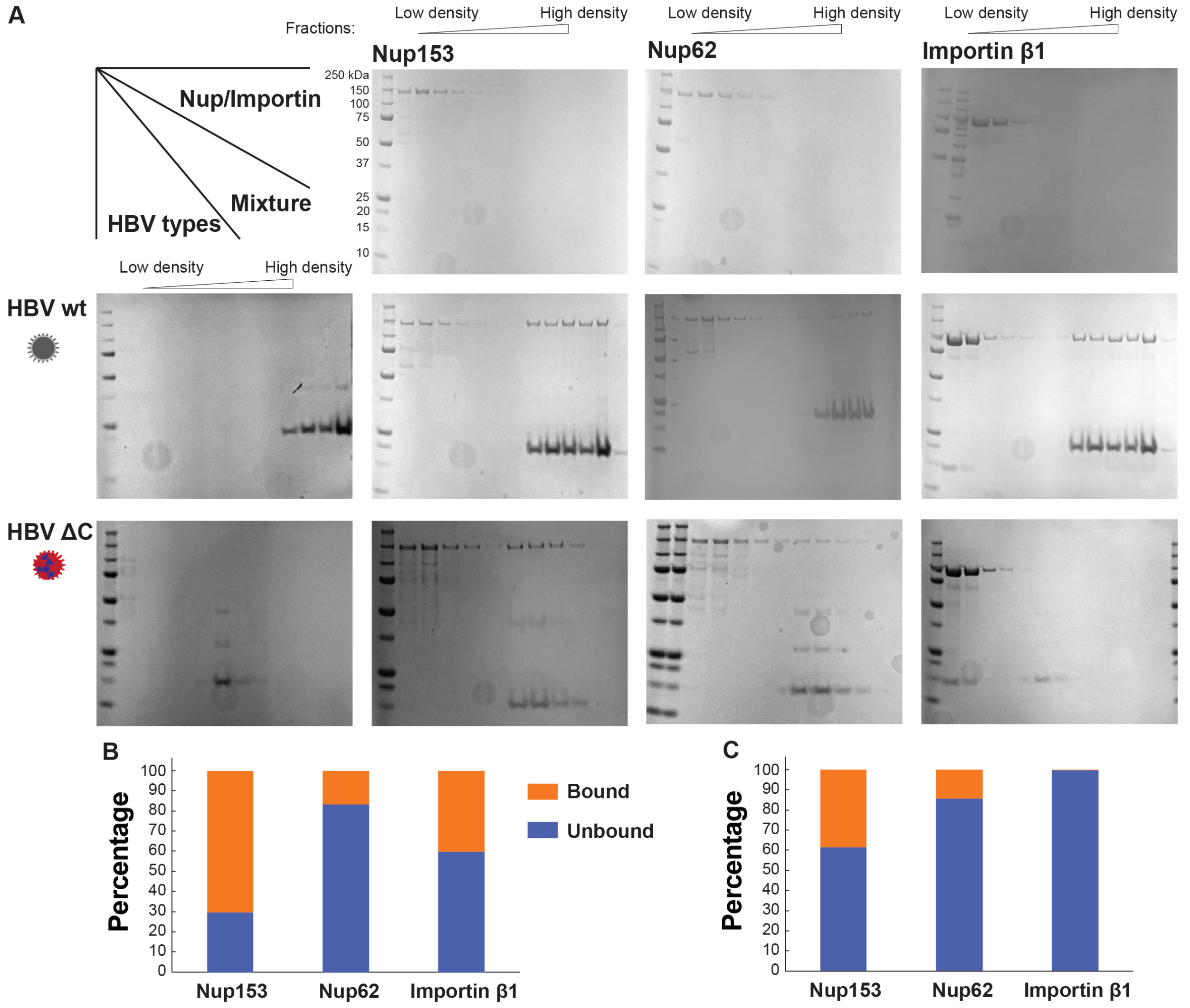


**Supplementary Figure 9**: Comigration assays showing HBV capsid binding with nups and importin β1. **(A)** SDS-PAGE characterization of the fractions recovered from the comigration assay, in which nups or importin (1 µM) were mixed with capsids (~3 µM of HBc), loaded to the top of 15%–45% glycerol gradients, and spun at 48,000 rpm for 90 min. Lighter fractions are shown on the left. Gels in the top row contain pure nups and importins. Gels in the left column contain purified HBV capsids. The rest of gels contain capsids mixed with nup or importin, the contents of which are identified by the row/column headers. Capsid binding is indicated by the existence of nup or importin in the 4–5 heaviest fractions where capsids reside. **(B)** Band intensities measured from gels characterizing nup153 (blue), nup62 (orange) and importin β1 (grey) binding to the wild-type (wt) HBV capsid, normalized to the total band intensity of the respective proteins across the whole gradient. **(C)** Same as (B), except for the HBV capsid with C-terminus deletion (ΔC).

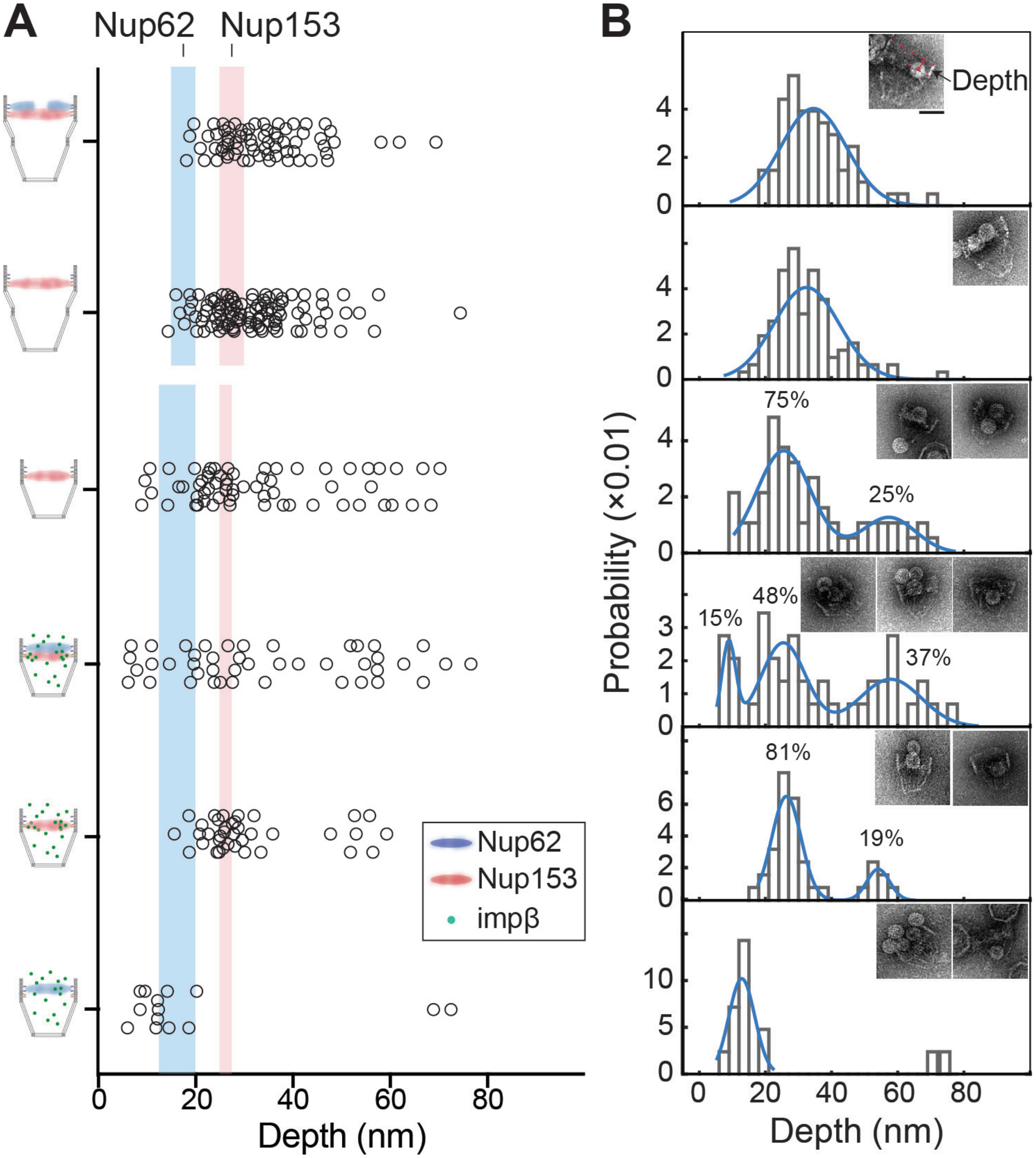


**Supplementary Figure 10**: Depths of HBV capsid penetration into NuPODs.

**(A)** Scatter plot of the capsid penetration depths (from the center of capsids to the opening of NuPODs). Blue and red bands denote the anchoring positions of Nup62 and Nup153 in NuPODs, respectively. NuPOD schematics are shown on the left of the scatter plot. When presented, importin β1 (impβ) concentration is 100 nM. **(B)** The capsid penetration depths plotted as histograms with Gaussian fit. When fitted using multiple Gaussian curves, the percentage and a representative TEM image of each population are shown. Scale bar: 50 nm. Number of NuPODs measured: 79-nm Nup62-Nup153 NuPODs (n=68); 79-nm Nup153 NuPODs (n=104); 60-nm Nup153 NuPODs (n=62); 60-nm Nup62-Nup153 NuPODs with impβ (n=43); 60-nm nup153 NuPODs with impβ (n=37); 60-nm Nup62 NuPODs with impβ (n=14).

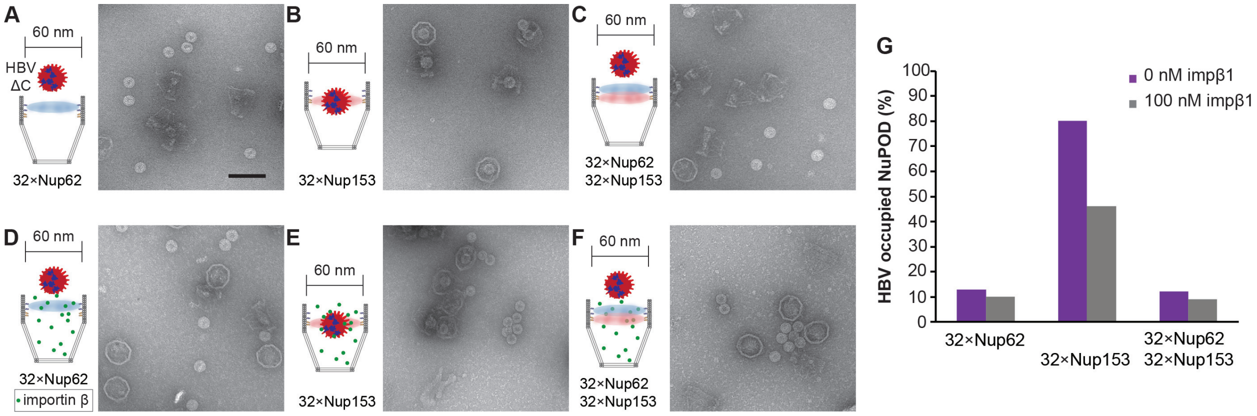


**Supplementary Figure 11**: HBV capsids with C-terminus deletion ($\Delta$C) interacting with NuPODs. **(A)** HBV$^{\Delta C}$ capsids mixed with capped 60-nm Nup62 NuPODs. **(B)** HBV$^{\Delta C}$ capsids mixed with capped 60-nm Nup153 NuPODs. **(C)** HBV$^{\Delta C}$ capsids mixed with capped 60-nm Nup62-Nup153 NuPODs. **(D)**–**(F)** are the same as **(A)**–**(C)**, but in the presence of 100 nM importin $\beta$1. For **(A)**–**(F)**, schematic diagrams of the binding experiments are shown next to representative TEM images. Scale bar: 100 nm. **(G)** Percentages of NuPODs occupied by the HBV$^{\Delta C}$ capsids in experiments **(A)**–**(F)**. NuPODs counted in each experiment are (from left to right): 131, 299, 252, 454, 250 and 201.

**Supplementary Movie 1**: AFM recording of 60-nm Nup62 NuPOD in the presence of 100 nM importin $\beta$1. Scale bar: 10 nm.

**Supplementary Movie 2**: AFM recording of 79-nm Nup62 NuPOD in the presence of 100 nM importin $\beta$1. Scale bar: 10 nm.
